# Environment and diet shape the geography-specific *Drosophila melanogaster* microbiota composition

**DOI:** 10.1101/2024.10.07.617096

**Authors:** Joseph T. Gale, Rebecca Kreutz, Sarah J. Gottfredson Morgan, Emma K. Davis, Connor Hough, Wendy A. Cisneros Cancino, Brittany Burnside, Ryan Barney, Reese Hunsaker, Ashton Tanner Hoyt, Aubrey Cluff, Maggie Nosker, Chandler Sefcik, Eliza Beales, Jack Beltz, Paul B. Frandsen, Paul Schmidt, John M. Chaston

**Affiliations:** Department of Plant and Wildlife Sciences, Brigham Young University, Provo, Utah, USA, 84602; Department of Biology, University of Pennsylvania, Philadelphia, PA 19104

**Keywords:** latitude, temperature, photoperiod, neutral theory, lactic acid bacteria, local adaptation, life history, rapid adaptation

## Abstract

Geographic and environmental variation in the animal microbiota can be directly linked to the evolution and wild fitness of their hosts but often appears to be disordered. Here, we sought to better understand patterns that underlie wild variation in the microbiota composition of *Drosophila melanogaster*. First, environmental temperature predicted geographic variation in fly microbial communities better than latitude did. The microbiota also differed between wild flies and their diets, supporting previous conclusions that the fly microbiota is not merely a reflection of diet. Flies feeding on different diets varied significantly in their microbiota composition, and flies sampled from individual apples were exceptionally depauperate for the Lactic Acid Bacteria (LAB), a major bacterial group in wild and laboratory flies. However, flies bore significantly more LAB when sampled from other fruits or compost piles. Follow-up analyses revealed that LAB abundance in the flies uniquely responds to fruit decomposition, whereas other microbiota members better indicate temporal seasonal progression. Finally, we show that diet-dependent variation in the fly microbiota is associated with phenotypic differentiation of fly lines collected in a single orchard. These last findings link covariation between the flies’ dietary history, microbiota composition, and genetic variation across relatively small (single-orchard) landscapes, reinforcing the critical role that environment-dependent variation in microbiota composition can play in local adaptation and genomic differentiation of a model animal host.

**SIGNIFICANCE STATEMENT:** The microbial communities of animals influence their hosts’ evolution and wild fitness, but it is hard to predict and explain how the microbiota varies in wild animals. Here, we describe that the microbiota composition of wild *Drosophila melanogaster* can be ordered by temperature, humidity, geographic distance, diet decomposition, and diet type. We show how these determinants of microbiota variation can help explain lactic acid bacteria (LAB) abundance in the flies, including the rarity of LAB in some previous studies. Finally, we show that wild fly phenotypes segregate with the flies’ diet and microbiota composition, illuminating links between the microbiota and host evolution. Together, these findings help explain how variation in microbiota compositions can shape an animal’s life history.

## INTRODUCTION

Animal-associated microorganisms (‘microbiota’) can profoundly impact the behavior, physiology, and evolution of their hosts. The types and abundances of microorganisms in wild animals, especially animals colonized by horizontally-acquired poly-species communities, often vary dramatically in response to numerous factors, including diet, time, and space. Because the types and abundances of the microorganisms can determine specific host traits, these changes in microbiota composition can influence adaptation of their hosts (1). Here, we sought to better understand the causes of variation in the microbiota composition of animals in the wild by studying the microbiota of the fruit fly *Drosophila melanogaster*, a host with a relatively low-diversity and low-abundance microbial community that is a model for understanding patterns of microbial community assembly (2), microbe-microbe interactions (3, 4), and host-microbe interactions, including in a wild setting (1, 5–7).

As in many other animals with a horizontally-acquired gut microbiota, the *D. melanogaster* microbiota composition is driven by the diet, the host, and other members of the community. In the wild or the laboratory, a single fly typically bears several hundred thousand bacterial cells from fewer than 100 species, most of which are consolidated in fewer than 10 highly abundant species (8–17). Flies are typically dominated by Acetic Acid Bacteria (AAB), Lactic Acid Bacteria (LAB), or bacteria from the order Enterobacteriaceae (5, 6, 8–10, 12, 15, 18–20), and the types and abundances of these microorganisms are typically influenced by the same factors as influence the gut microbiota in mammals: diet (15, 16, 20), host genotype (21), and individuality, including vial effects for flies reared in the same containers (16, 22). In the laboratory, members of the *Acetobacter*, *Lentilactobacillus*, and *Lactoplantibacillus* genera have been commonly used as representative isolates in the flies (23). Studies with bacteria from these and other genera have revealed that some, but not all bacterial strains persistently colonize the flies for longer than the bulk passage of diet through the gut and that there are specific foregut niches for some of these persistently colonizing bacteria (22, 24–28). Also, the types and abundances of the microbes in the flies are distinct from those in the diet (16), dramatically influenced by host genetic selection and microbe-microbe interactions (3, 21, 23, 29, 30), and typically best defined by neutral community assembly rules, suggesting strong roles for ecological drift and passive dispersal (6, 12). Thus, a substantial body of work has established key drivers of microbial community structure.

The fruit fly *D. melanogaster* is an established model for studying geographic variation and the microbiota. The geographic life history of *D. melanogaster* may be the most extensively studied of any animal on the planet, with hundreds of studies documenting latitudinal clines in allele frequencies at candidate genes (e.g. (31–35)) and fitness-associated traits (e.g. (36–40)), and in patterns of genomic differentiation (e.g. (41–43)). Populations of *D. melanogaster* in the eastern United States show clear genomic and phenotypic differentiation between different latitudes and seasons. The microbiota composition of *D. melanogaster* also varies across these geographic and seasonal clines, and can profoundly influence the life history traits and evolution of its host (1, 5, 6). Two recent studies of the microbiota of wild flies from the eastern USA have each reported substantial geographic variation in microbiota composition, with some conflicting findings between the studies. In one of these studies, we reported substantial numbers of LAB in many of the sampled flies, a latitudinal gradient in the AAB:LAB ratio, and suggested that this pattern revealed congruence between microbial abundance, the influence of those microorganisms on host traits, and the host traits naturally adopted in the sampled locations (5). Another study sampled deeply at more locations and found essentially no LAB in any of the samples and little evidence for a latitudinal cline within- or between-host variation in microbiota composition (6). The authors concluded that variation in microbiota composition was determined primarily by neutral processes and strict host filtering. Also, following evidence that LAB are rare in some (6, 15) but not other (5, 12, 18–20) samplings of wild flies, the authors suggested that LAB may be more likely to colonize laboratory than wild flies. These disparate findings show gaps in our understanding of what determines the wild fly microbiota composition.

To better understand the relationship between sampling location and *D. melanogaster* microbiota composition, we asked four major questions: 1) Can previously observed latitudinal patterns in microbiota composition be observed in fresh samplings of wild flies? 2) What is the relationship between the microbiota of wild flies and their wild diet? 3) Why are LAB readily recovered in some, but not all wild *D. melanogaster* samplings? 4) How is variation in the microbiota composition of wild flies related to their life history? We addressed these questions by comparing the sequencing results of previous and new collections of wild *D. melanogaster*, measuring the microbiota composition of wild and laboratory fly populations reared on distinct environmental conditions or on different diets, and experimentally dissecting the contributions of time and diet decomposition to microbiota composition. We also measured a key life history trait in wild-caught, laboratory-reared fly populations. Together, these results evidence that specific environmental conditions predict patterns in microbiota composition better than latitude, that diet influences but does not necessarily seed the wild fly microbiota, and that fly phenotypes can segregate with the diet of flies in the wild.

## RESULTS

### A geography-specific *D. melanogaster* microbiota composition is associated with environmental temperature

We reanalyzed two previously published samplings of flies in the eastern USA and compared these to the results of freshly collected samples, to better understand the reasons for their different outcomes. The previous studies included samples collected in 2009 (**Fig. 1B**, from apples and peaches) and 2018 (**Fig. 1C** (from grapes)-**D** (from apples)). We added two additional samplings, one in the eastern USA (**Fig. 1E**, 2021, from apples) and one in the state of Utah, USA (**Fig. S1**, 2020, from peaches). Each of the eastern USA collections included at least some sites shared with the other studies. The most abundant genera in the new samplings generally mirrored the previous studies, including that the flies were dominated especially by AAB, with substantial numbers of Enterobacteria or LAB in certain samples. Mantel tests of the relationship between microbiota composition and latitude recapitulated previous conclusions by showing that the 2009, but not the 2018, microbiota composition significantly covaried with latitudinal distance (**Fig. 1F-H, Fig. S2**). As in 2009, the microbiota composition of the two new samplings covaried with latitude (**Fig. 1I**, **Fig. S1C**). We sought to reconcile these different outcomes by testing if environmental conditions could help expain the variation in microbiota composition from different locations. Of 41 variables we tested, the microbiota in all five locations significantly covaried with just one factor - the daily maximum temperature (**TABLE S3**) - suggesting temperature could be a major determinant of wild fly microbiota composition (**Fig. 1J-M, Fig. S1D**).

**Figure 1.**
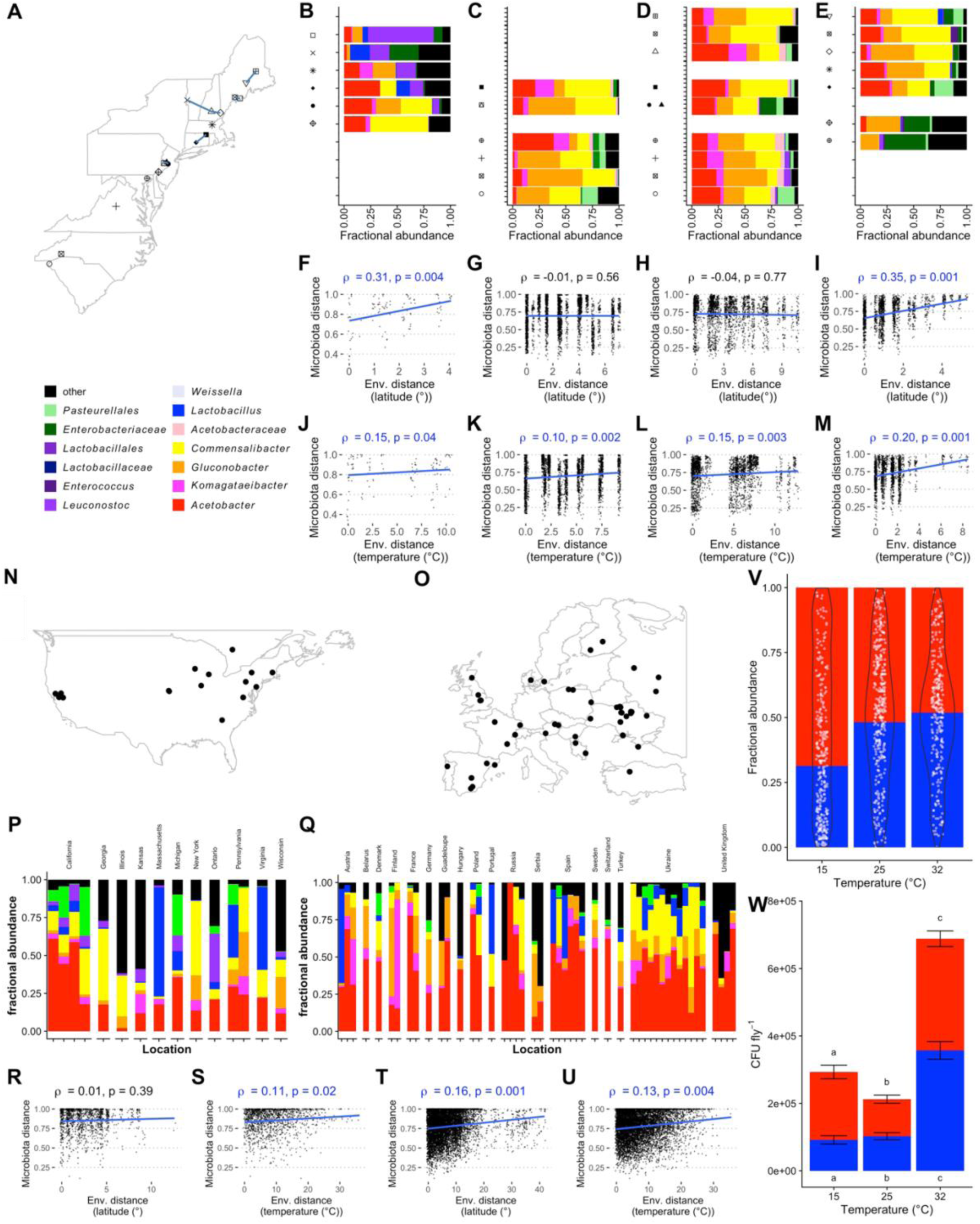
Latitude- and temperature-dependent variation in microbiota composition of *D. melanogaster* from the eastern USA in multiple years and samplings. *Drosophila melanogaster* from (**A**) the same general area in the eastern USA were sampled and sequenced across portions of the 16S rRNA V4 regions by two laboratories in **B**) 2009 (apples and peaches, N=15), **C**) 2018 (grapes, N=71); **D**) 2018 (apples, N=79), and **E**) 2021 (apples, N=60). Taxon plots are an average of samples which are pools of flies (**B**, rarefaction threshold (rt) = 475) or individual flies (**C-E**, rarefaction threshold= 500). Mantel tests reporting the relationship of the Bray Curtis and environmental distances between each sample were also performed, with environmental distance calculated from distance matrices based on **F-I**) latitude or **J-M**) latitude plus maximum temperature and minimum relative humidity (RH) on the day of sampling. Letters over **F-M** report the Mantel test correlation coefficient ρ and the p-value results when the results are (blue) or are not (black) significant. **P-Q**) A similar analysis as for eastern USA flies, conducted using sequences extracted from whole genome sequencing of flies collected with the DEST dataset. **N-O**) Maps, **P-Q**) taxon plots, and **R-U**) Mantel test results for DEST flies from N,P,R,S) North American and **O,Q,T,U**) Europe are reported. Samplings from Guadeloupe, a department of France in the Caribbean, is not shown on the map. **V-W**) Common garden populations of individual isofemale lines from each location in **F_IG_. 2** were reared under 6-species gnotobiotic conditions in the laboratory until adult flies were 3 days old, transferred to test conditions for 3 days, then the microbiota composition of pools of 2 surface-sterilized adults was analyzed by homogenization and dilution plating (2 pools of 2 females and 2 males per vial, 3 vials per common garden population in each experiment, 3 separate experiments in time). The relative **V**) and absolute **W**) abundances of AAB (red) and LAB (blue) colony forming units (CFUs) in flies reared at varying temperatures. Relative abundances are shown as the mean of AAB counts divided by the mean of LAB counts, with the fraction of LAB shown as a white point and the overlayed violin plot showing the distribution of fractional LAB abundance. Significant differences in relative abundances of LAB were determined by PERMANOVA (**T_ABLE_ S4**). Absolute CFU abundances are shown as the mean and standard error of the mean of all replicates. Significant differences in AAB and LAB abundance were determined by a Kruskal-Wallis test with a post-hoc Dunn test, and different letters over (AAB) or under (LAB) the bars report significant differences in their abundance

We then tested if these findings applied to flies sampled in other areas and at various times by comparing the same environmental and geographic metrics with the microbiota composition of flies collected as part of the Drosophila Evolution over Space and Time (DEST) data set, which were collected across North America and western Europe mostly between 2014 and 2016 (**Fig. 1N-Q**, (44)). As before, the geographic and microbiota distances covaried significantly in one, but not both sampling areas (**Fig. 1R,T**). However, the difference in maximum temperature on the day of sampling was significantly correlated with the distance in microbiota composition in samples from both North America and Europe (**Fig. 1S,U**, **TABLE S3**). We also confirmed experimentally that variation in temperature significantly determines gnotobiotic microbiota composition (**Fig. 1V-W**, **TABLE S4**, **Fig. S3**). Beyond temperature, UV irradiance and humidity also covaried with microbiota composition in many samplings (**TABLE S3**). We investigated microbiota effects of humidity previously (45), and report here that photoperiod is also associated with significant changes in microbiota composition, though the changes are less marked than those caused by temperature (**Fig. S4, TABLE S4**). From these findings temperature emerges as a primary, but not the only, driver of variation in wild *D. melanogaster* microbiota composition.

### The *D. melanogaster* microbiota is not a reflection of the microbiota in its diet

To further define the contributions of a wild environment to wild flies’ microbiota, we compared community composition in flies bearing a total (resident + transient) microbial community, flies bearing only a resident microbiota, and neighboring fruits and soils. We report on comparisons between individual fruits, wild flies captured from those fruits, and adjacent soil samples. Low prevalence of *D. melanogaster* in some sites meant that we did not obtain samples of flies bearing a total and resident-only microbial community from all fruit samplings, but among the fruit sites that did have *D. melanogaster* we sequenced their microbiota and compared it to the corresponding fruit and sample locations (**Fig. 2A-B**). These samples’ microbiota composition varied significantly with both the sample type (resident fly microbiota, fruit, etc, F_3,218_ = 20.08, R^2^ = 0.20, p < 10^-4^, **TABLE S5**) and the orchard they were sampled from (F_7,218_ = 2.28, R^2^ = 0.05, p < 10^-4^, **Table S5**). No bacterial genera varied significantly in abundance between the total and resident portion of the microbiota, but there were significant differences in the abundances of bacteria between the diet and wild flies, consistent with the expectation of strict filtering by the flies (6). More than 85% of total reads were assigned to the < 20% of ASVs that were shared between fruit and flies (**Fig. 2C**, **Table 1**), and among these *Commensalibacter*, *Lactobacillus*, *Enterococcus* and reads that could not be assigned below the *Pasteurellales* level were all significantly more abundant in flies than diets (**Table 1**). Other bacterial strains were more abundant in the diet than the flies (**Table 1**), together identifying differences between the microbiota of flies and their diets and classifying bacteria from the different genera as fly- or diet-preferred. Across both types of fly samples, neutral modeling reported lower AIC scores when microbiota was sampled from itself rather than diet, further confirming that the patterns in microbiota composition follow neutral patterns ((6), **Fig. 2D**). Also, the observation that the beta-diversity distances between fruits and the total or resident microbiota were similar tends to confirm the conclusion that the colonization niche of the bacteria does not influence similarity to the diet (**Fig. 2E**).

**Figure 2.**
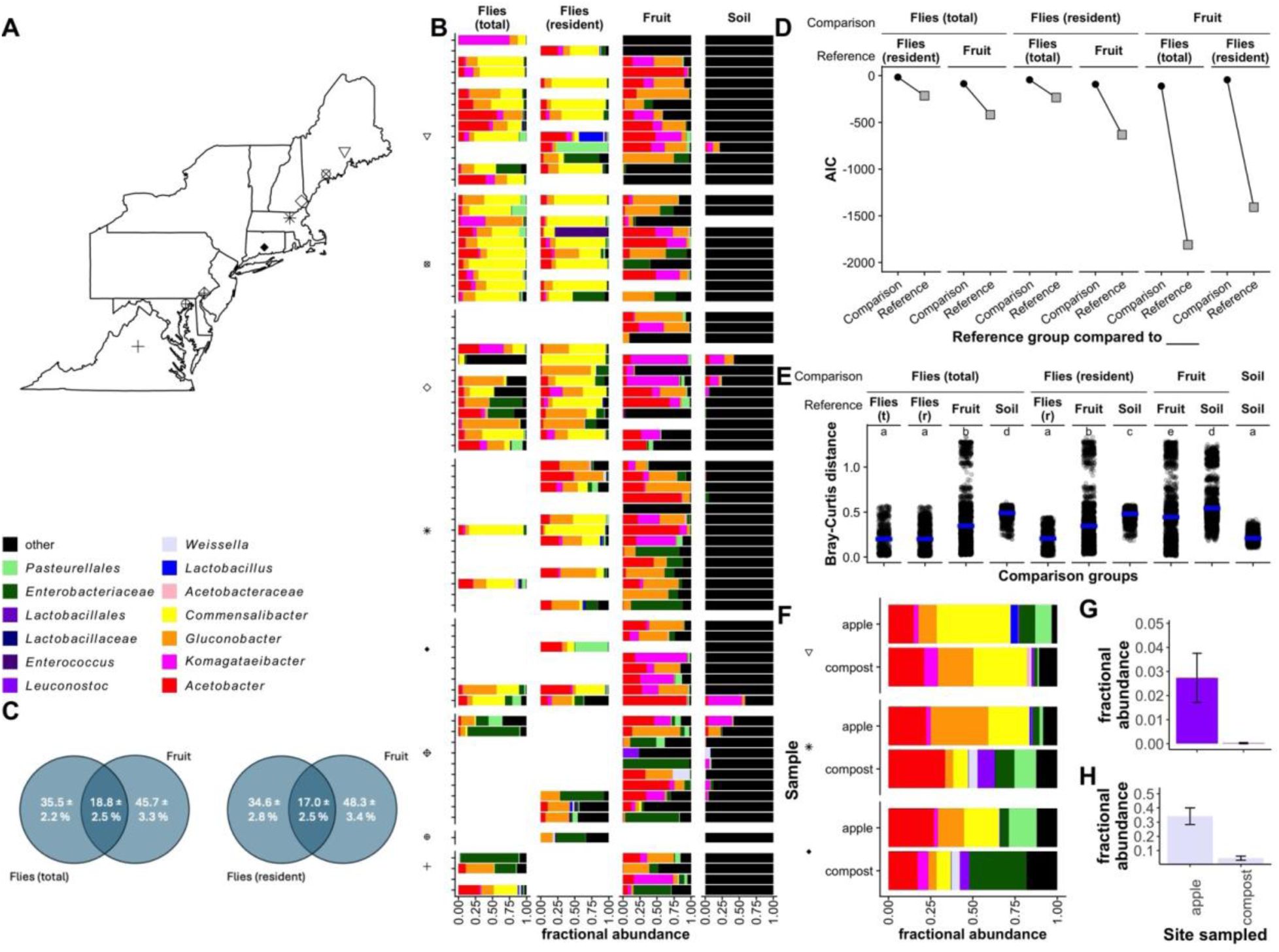
The total and resident *D. melanogaster* microbiota composition is distinct from the diets of wild flies. A-B) The 16S rRNA V4 region was sequenced in samples of individual wild *Drosophila melanogaster*, their diets, and nearby soil locations from multiple locations in the eastern USA. Flies were immediately frozen after collection (‘Flies (total)’) or starved in empty vials for > 2 h (‘Flies (resident)’) after transient microorganisms had passed through the fly gut with the bulk flow of diet. C-D) Reads assigned to the genus *Commensalibacter* are shown in different samples. * = significant differences in abundance by ANCOM. E) Venn diagrams showing the average fractional abundance of ASVs ± s.e.m. that were unique to or shared between sampled fruit and the total or resident fly microbiota. F) AIC values for neutral models calculated with the group indicated in the ‘Reference’ row was compared to itself (‘Reference’, gray squares) or the group in the ‘Comparison’ (black squares) row. G) Weighted Unifrac distances between samples assigned to the groups in the Reference and Comparison rows, including the mean distance between samples (blue bar). Different letters over the clusters of points represent statistically significant differences in distance between the comparison groups as determined by a Kruskal-Wallis test with a post-hoc Dunn test. F-G) Microbiota composition was measured In individual apples or compost piles at multiple locations in the eastern USA, Bars are the averages of mulitiple samples (N = mean 6.5 ± sem 1.2, min = 3, max = 10 samples per bar condition) rarefied to 845 reads each. Reads assigned to the genera E) *Leuconostoc* and F) *Weissella* differed significantly in abundance between locations as determined by ANCOM.

**Table 1.** Read counts of bacterial genera in flies relative to the fruits they were feeding on.

| Genus | Flies (total microbiota) |  |  |  | Flies (resident microbiota) |  |  |  |
| --- | --- | --- | --- | --- | --- | --- | --- | --- |
|  | Flies | Fruit | % in fly | p-value | Flies | Fruit | % in fly | chisq_p |
| *Commensalibacter* | 6113 | 0 | 100 | $10^{-99}$ | 6675 | 23 | 100 | $10^{-99}$ |
| *Lactobacillus* | | | n.t. | | 255 | 2 | 99 | $10^{-54}$ |
| *Enterococcus* | | | n.t. | | 416 | 18 | 96 | $10^{-79}$ |
| Unassigned<br>*Enterobacteriaceae* | 2009 | 1187 | 63 | $10^{-43}$ | 1474 | 1683 | 47 | 0.0004 |
| Unassigned *Pasteurellales* | 641 | 393 | 62 | $10^{-13}$ | 550 | 290 | 66 | $10^{-17}$ |
| *Pseudomonas* | 92 | 100 | 48 | 0.62 | 61 | 189 | 24 | $10^{-14}$ |
| *Gluconobacter* | 3484 | 3878 | 47 | 0.00004 | 3839 | 3680 | 51 | 0.10 |
| *Acetobacter* | 2418 | 4565 | 35 | $10^{-121}$ | 2342 | 3923 | 37 | $10^{-74}$ |
| *Gluconacetobacter* | 1120 | 2574 | 30 | $10^{-113}$ | 489 | 2932 | 14 | $10^{-99}$ |
| Unassigned<br>*Xanthomonadaceae* | 62 | 193 | 24 | $10^{-15}$ | 91 | 213 | 30 | $10^{-11}$ |
| Unassigned *Bacteria* | 88 | 1420 | 6 | $10^{-246}$ | 59 | 1991 | 3 | $10^{-99}$ |
| *Lactococcus* | 5 | 274 | 2 | $10^{-56}$ | 9 | 356 | 2 | $10^{-71}$ |
| Unassigned *Proteobacteria* | 1 | 393 | 0 | $10^{-84}$ | | | n.t. | |

<sup>a</sup> n.t. = not tested, < 1% of total reads collected

Separately, the resident microbiota of flies collected from compost piles at three of these sites was significantly different from the composition in flies from fallen apples (F_1, 31_ = 2.11, R^2^ = 0.05, p = 0.01, **TABLE S6**, **Fig. 2F**; these were the only sites with compost piles, samplings from compost and apples were done at the same time, and the apple bars include data shown in **Fig. 2B**). This finding is somewhat surprising because we assumed microbiota composition would be relatively homogenous across an orchard as *Drosophila* can readily migrate miles overnight (46–48). Two genera differed in abundance in flies sampled from apples and compost piles, and both were members of the LAB: *Leuconostoc* and *Weissella* (**Fig. 2G-H**). These findings confirm that there is diet-dependent variation in microbiota composition of flies within the same orchard. The data also reveal that flies sampled from apples are less likely to bear LAB abundantly, and may help to explain why two recent collections of flies from individual fallen apples reported low levels of LAB (**Fig. 1C-E**). However, because the compost piles had many different types of decomposing material and were not just apples, it is not clear if the differences in fly microbiota composition when collected from individual fruits or composted material are due to differences in the substrate or to its decomposition state.

Thus, while diet contributes to variation in microbiota composition, both the resident and transient portions of the microbiota can be distinct from their proximal dietary sources at the time of collection. The differences may be due to filtering, as has been asserted previously (6), but the very low abundance and prevalence of the fly-preferred microbes in the flies’ diets also suggests transmission fidelity between flies. Alternatively, the flies may have obtained their microorganisms from diets distinct from those that they were feeding on at time of capture, although the finding that the microbiota differs between flies feeding on different diets in the same orchard is inconsistent with this expectation.

### Wild diets determine differences in the *D. melanogaster* microbiota composition

To follow up on the role of diet in shaping the fly microbiota composition, especially the abundance of LAB, we compared the microbiota composition of flies sampled from different wild diets in a single orchard. The flies’ microbiota significantly varied when sampled individually from varieties of apple, peach, and pear in a single orchard (F_2, 31_ = 2.04, R^2^ = 0.10, p < 10^-4^, **TABLE S7**). However, no bacterial groups at any taxonomic level varied significantly with the type of fruit the flies were sampled from. We adopted a more controlled approach by showing that microbiota composition in laboratory flies reared under gnotobiotic conditions with 12 different bacterial species varied significantly with the type of fruit the flies were reared on (F_9, 52_ = 2.42, R^2^ = 0.34, p < 10^-4^, **TABLE S8**). Four of the species varied significantly in abundance in flies reared on the different fruits: *L. brevis, W. paramesenteroides*, *G. cerinus,* and *Pantoea* sp. JGM106 (ANCOM W score = 11 for each species)*. L. brevis* was particularly prevalent in peaches, pears, and oranges, and LAB also tended to have higher average abundance in the non-apple samples from our wild samplings (**Fig. 3A**). Additionally, apples had far greater average AAB abundance than any other samples, consistent with observations that flies sampled from apples were dominated by AAB (**Fig. 1C-E**), and that our samplings in Utah, which were from peaches, frequently bore abundant LAB (**Fig. S1**). These findings reveal that the *D. melanogaster* microbiota varies with diet in both the wild and laboratory, including in flies reared with the same starting set of microorganisms.

**Figure 3.**
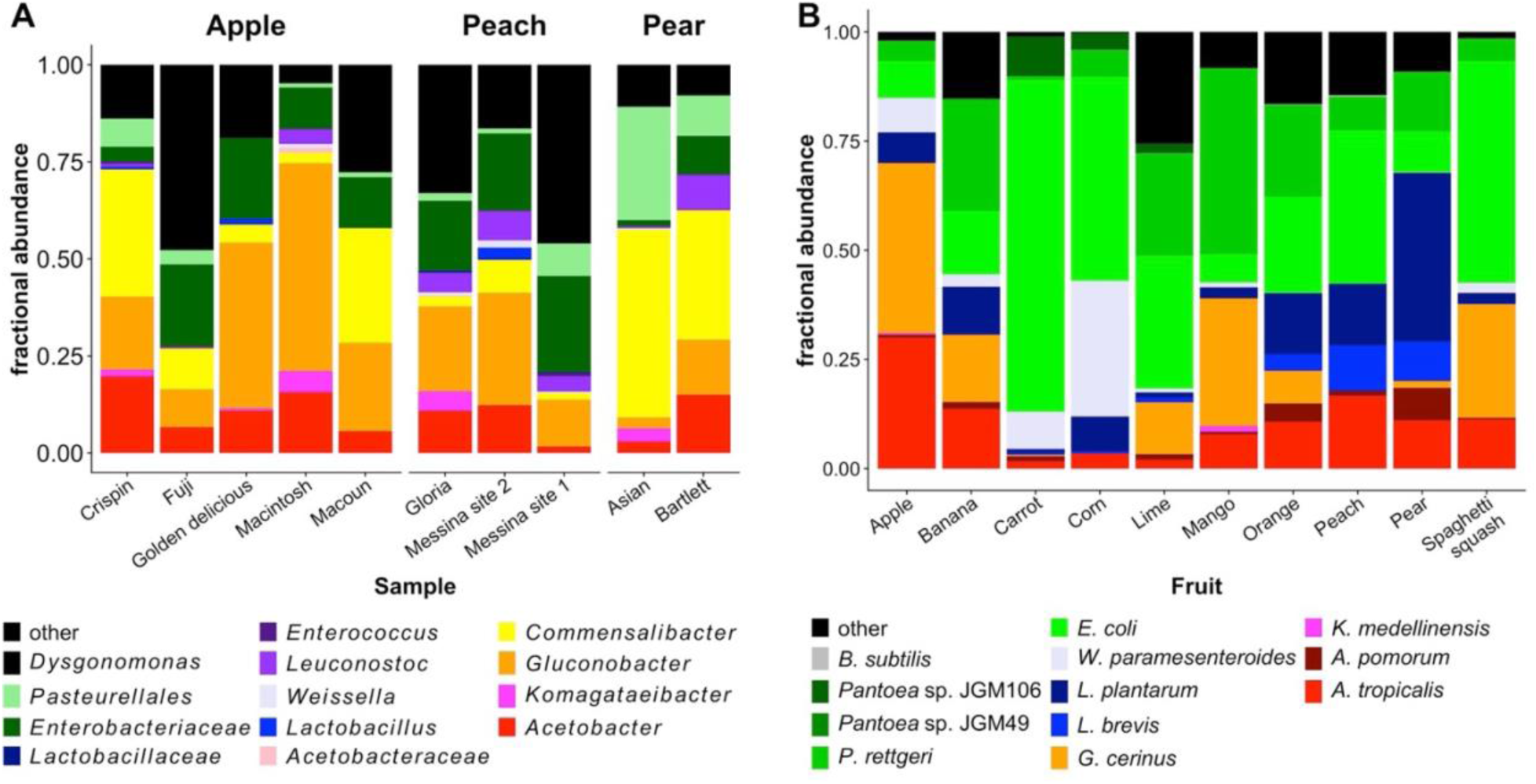
The microbiota composition of wild and laboratory *D. melanogaster* varies with diet type. We sequenced the 16S rRNA V4 region of the *D. melanogaster* microbiota collected from different fruits in the wild and the laboratory. A) Composition of the resident microbiota in wild male *D. melanogaster* collected from fallen fruit at Lyman Orchards in Middlefield, CT. Bars are the averages of mulitiple samples each rarefied to 120 reads (N = mean 3.2 ± sem 0.25, min = 2, max = 4 samples per bar). B) Composition of the total microbiota in gnotobiotic 12-sp *D. melanogaster* CantonS male flies. Bars are the averages of mulitiple samples each rarefied to 200 reads (N = mean 5.3 ± sem 0.54, min = 3, max = 7 samples per bar).

### Population dynamics of the wild *Drosophila* microbiota are distinct with time and decomposition of wild fly diets

We next investigated how time and diet decomposition shape the microbiota composition of wild flies, especially their LAB abundance, by capturing flies from fruit piles in different decomposition states. In a single fall season, we established separate piles of peaches and apples every two weeks and sequenced the resident microbiota of flies sampled twice weekly from the various piles. The microbiota of the flies varied with the type of fruit they were sampled from (F_1, 183_ = 11.79, R^2^ = 0.05, p < 10^-4^, **TABLE S9**) and over time (F_1, 183_ = 10.5, R^2^ = 0.05, p < 10^-4^, **TABLE S9**) (**Fig. 4A**). Also, the overall microbiota composition covaried with time (**Fig. 4B**, **Fig. S5**, s = 0.12, p < 10^-4^). *Gluconacetobacter*, *Leuconostoc*, and reads that could not be assigned below the family Enterobacteriaceae significantly varied over time, and each was significantly correlated with time (**Fig. 4D-E**). No genera varied significantly in flies sampled only from apples, but *Gluconacetobacter* and *Lactobacillus* reads varied significantly with time in flies sampled from peaches (**Fig. 4F-G**).

**Figure 4.**
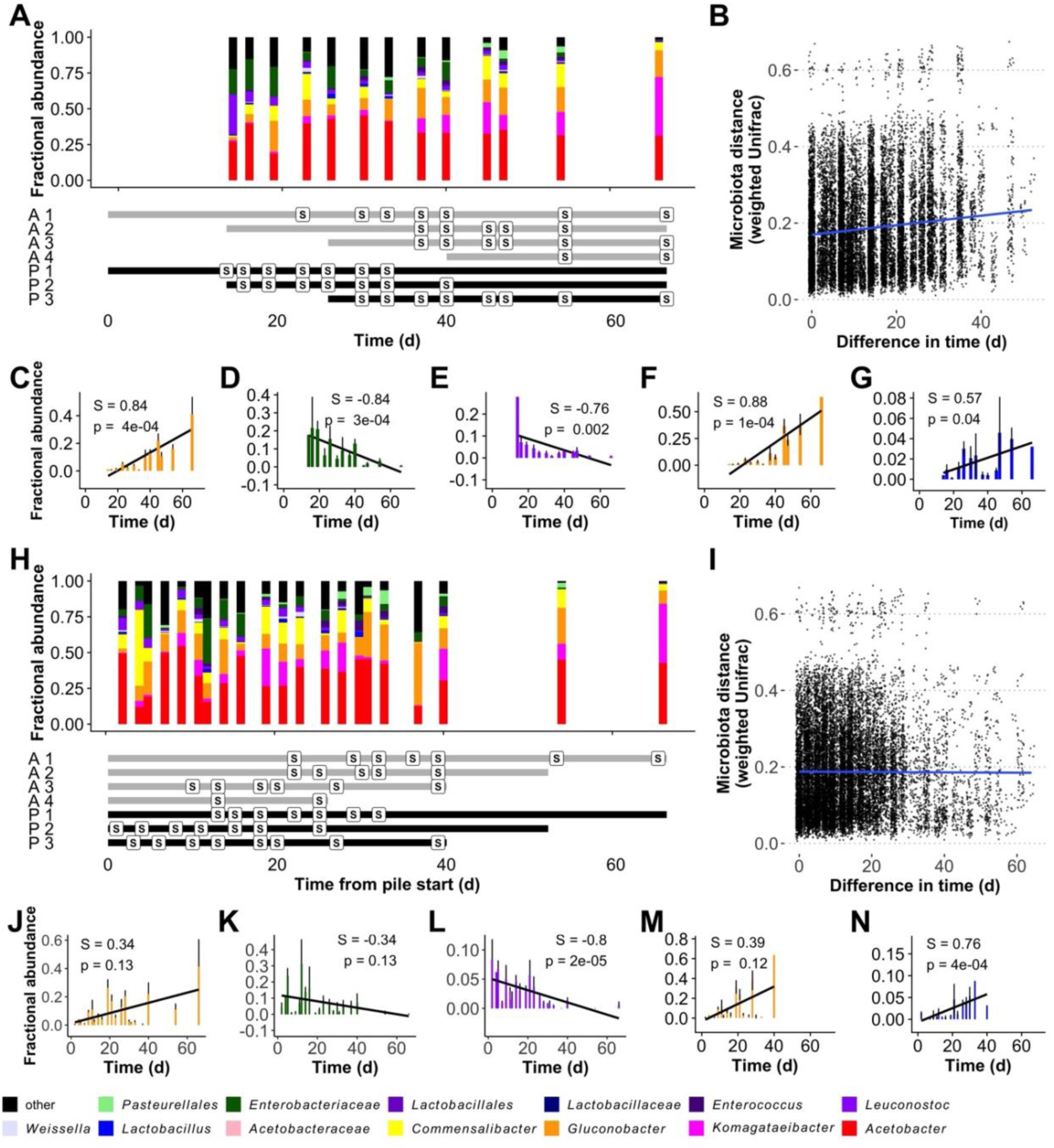
The microbiota composition of wild *D. melanogaster* varies with time and diet decomposition. We sequenced the 16S rRNA V4 region of the resident microbiota in individual wild *Drosophila* collected from 1-bushel piles of fruit established at different times in an experimental orchard in Provo, UT. The same data are shown on two timescales, either relative to A-G) calendar date (time = 0 when the first piles were established), or H-N) pile establishment time (time = 0 for each pile when it was established). A,H) Taxon plot, plus a timeline showing the establishment and sampling times (S) from each of four apple (A) and three peach (P) piles. B,I) Mantel test showing the relationship of the weighted Unifrac and calendar date (B) or pile establishment time (I) distances between each sample, including the slope of the trendline (blue). Plots showing the abundances of specific members of the microbiota that varied in C-E,J-L) apples and peaches, or only in peaches F-G, M-N) in one or both timescales. The Spearman’s rank correlation coefficient (S) and p-value (p), plus a trendline for change in abundance over time (black line) are shown for each.

Then, we tested if diet decomposition shaped microbiota composition by comparing the microbiota between these same samples when time was defined relative to the ‘establishment time’ of the pile instead of calendar date. There was a significant effect of establishment time (**Fig. 4H**, F_1, 183_ = 4.06, R^2^ = 0.02, p = 0.01) on microbiota composition, but microbiota composition did not covary with establishment time (**Fig. 4I**, **Fig. S5**, s = - 0.005; p = 0.50). Further, the *Gluconacetobacter* (all samples, **Fig. 4I**; peaches only, **Fig. 4K**) and *Enterobacteriaceae* (**Fig. 4J**) reads that covaried with time did not covary with establishment time of the piles. Conversely, *Leuconostoc* abundance in peaches and apples (**Fig. 4L**) and *Lactobacillus* abundance in peaches only (**Fig. 4N**) varied more significantly with establishment time than calendar date. Together, these findings suggest that the relative abundance of most of the genera that vary in abundance in the flies, covary with time, but that the populations of the two LAB that varied across the different samplings responded more to establishment time of the piles, or their decomposition, than calendar date. The absence of these patterns when apples were considered alone is consistent with our previous observations that LAB are generally depauperate in apple samples and shows some diets are incompatible with the seasonal patterns we document here. Together, these experiments reveal that variation in the abundance of LAB in the flies is complex and determined at least in part by interactions between time, diet decomposition, and microbial compatibility with the fly diet.

### Diet determines microbiota-dependent genetic stratification of *D. melanogaster* in a single orchard

Finally, we tested if diet-dependent variation in microbiota composition was associated with variation in the life history of flies in a wild setting. We reasoned that since the microbiota is an agent of selection that can drive adaptation of its host (1), flies from diets with distinct microbiota composition might segregate phenotypically. When we measured the development time to adulthood for wild-caught isofemale lines reared in the laboratory under gnotobiotic and bacteria-free conditions, there were significant, non-interactive effects of the dietary source (Z_1, 4173_ = 8.04, p < 10^-15^) and rearing condition (Z_1, 4173_ = 7.81, p < 10^-14^) of the flies (**Fig. 5A-B**, nonsignificant interaction was Z_3_, _4173_ = -1.54, p = 0.12). The flies sampled from compost piles (ο), which support greater levels of LAB in the flies (**Fig. 2G**), developed to reproductive maturity more quickly than flies sampled from apples in multiple locations throughout the orchard (μ,Δ). This pattern, where the flies richer in LAB develop to reproductive maturity more quickly than flies depauperate for LAB, mirrors the pattern observed in flies that were evolved in the wild while being fed *L. brevis* or *A. tropicalis*, then reared bacteria-free in the laboratory (**Fig. 5C,** Z_3_, _1810_ = 3.73, p = 0.00019). Together, these findings support the conclusion that diet-dependent microbiota composition is associated with the genotypic differentiation in two distinct fly populations sampled from the wild.

**Figure 5.**
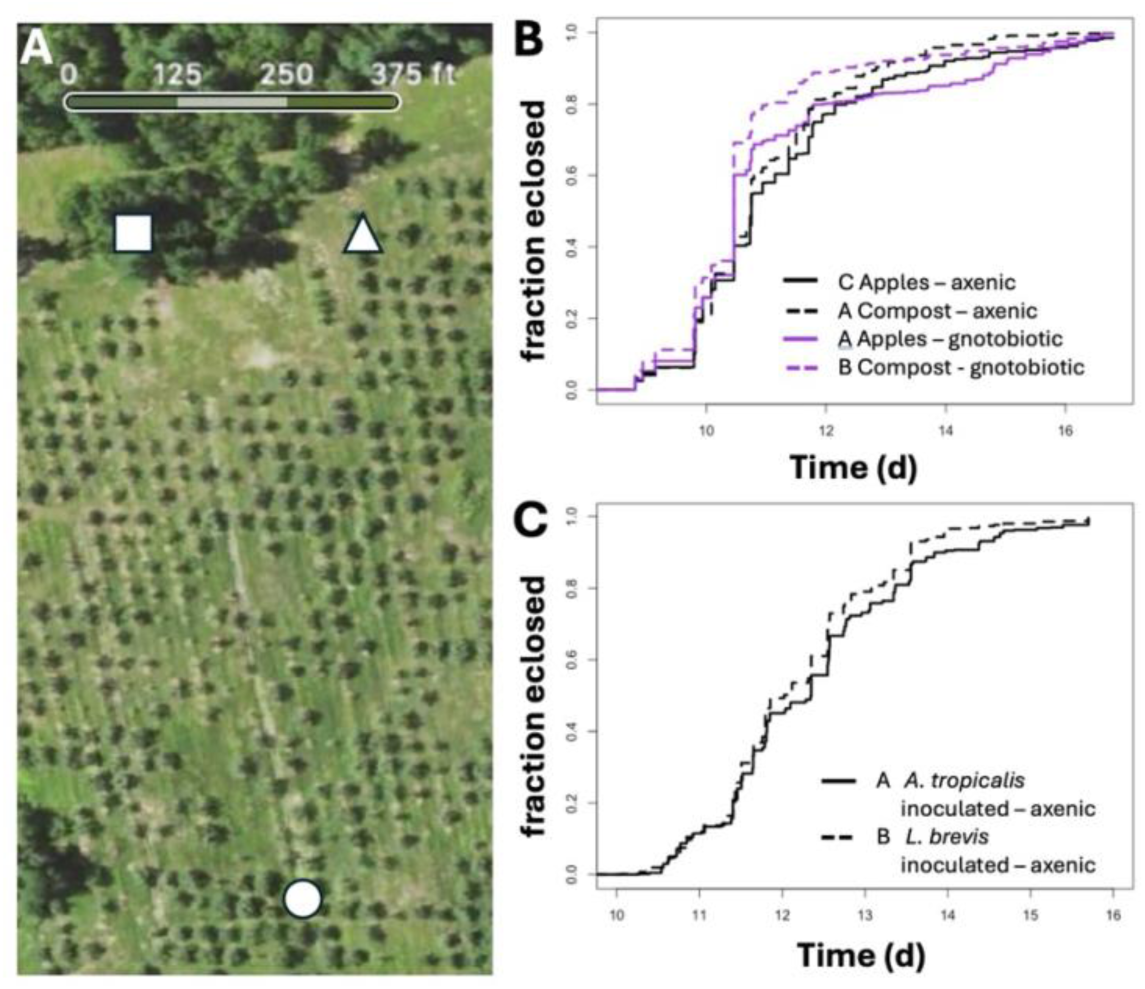
*D. melanogaster* phenotypes segregate with the diet they are collected from. A) Wild *D. melanogaster* were sampled from compost piles (ο) or individual fallen apples (μ,Δ) throughout a single orchard, shown in a screenshot from Apple maps taken Summer 2024. B) Isofemale lines derived from the wild collections were reared in the laboratory under gnotobiotic or bacteria-free conditions, and their development time to adulthood (eclosion) was measured. C) Development time to adulthood was measured in flies reported previously (1) when reared bacteria-free in the laboratory at the conclusion of selection. Significant differences between groups were determined by a Cox mixed-effect model.

## DISCUSSION

We sought here to define patterns in the microbiota composition of wild flies and reconcile differences between conflicting previous analyses. We found that latitudinal differentiation in microbiota composition could be better explained by considering temperature than latitude. Followup experiments in the laboratory confirmed that temperature and, to a lesser extent, photoperiod, influence microbiota composition (a role for humidity had been previously demonstrated (45)). Additionally, we provide evidence that even though the flies’ microbiota is not a direct reflection of their diet, the type and decomposition of fly diet influences distinct members of the fly microbiota. Whereas the AAB were prevalent and abundant in a variety of diets, LAB were most abundant in diets that were decomposing and were especially depauperate in flies sampled from apples. Together, these factors can help explain the varied differences in the microbiota composition of wild-caught flies, and show that numerous factors, most of which vary asynchronously with time and space, help determine variation in microbiota composition.

Diet can help explain some of the patterns in microbiota abundance observed across four different fly samplings (**Fig. 1**, **Fig. S1**). LAB were virtually absent in flies sampled from individual apples (**Fig. 1D-E**) and grapes (**Fig. 1C**), but were abundant when sampled from rotten peaches (**Fig. S1**) or not explicitly from individual fruits (**Fig. 1B**). We did not analyze the microbiota of gnotobiotic flies reared on grapes, but gnotobiotic flies reared on apples bore the highest fraction of AAB of any fruit we sampled, and, conversely, flies reared on peaches bore relatively high LAB loads. The wild samplings suggest that grapes, like apples, support relatively high AAB loads, and that the absence of LAB from the more recent east coast fly samples may have been driven by sampling from fruits that naturally support high levels of AAB (**Fig. 1C-E**). If samplings from individual apples or grapes, qualitatively assessed to be rotten, are less decomposed than a compost pile, then decomposition state may also contribute to these differences (but it is not possible to ascertain post-hoc). Diet type alone cannot explain differences in microbiota composition between experiments because **Fig. 1B** flies were sampled from apples (•, ∇, •) and peaches (ω,◊), and the peach-sampled flies had intermediate LAB abundance relative to the other apple samples. However, if the higher latitude samplings were from more decomposed fruits (not recorded at sampling), it could help explain why there were more LAB in flies collected from apples at high-latitudes than in peaches at lower latitudes. Regardless, these findings show that controlling for diet decomposition state and diet type in fruit flies provides context for variation in microbiota composition, and our approaches highlight one way that key drivers of microbiota variation can be identified in wild sampled animals.

Even though diet and its decomposition state were profoundly associated with variation in wild fly microbiota composition, the microbiota composition of wild flies and their proximal diets at time of collection were generally incongruent. The most abundant taxa in the flies were usually detected in the diet, and vice versa, but multiple approaches – comparison of ASV identities, modelling if fruit was the source of the fly microbiota, and directly comparing composition between sample types - all failed to link the abundance of those ASVs between the two sample types. The most conspicuous difference between the microbiota of flies and neighboring fruits was that *Commensalibacter* was the most abundant ASV in flies and essentially absent from the diet. *Lactobacillus* and *Enterococcus* were also abundant in flies but not their diets. These findings are consistent with the substantial body of work showing that laboratory and wild flies can be colonized persistently by specific sets of microorganisms (22, 24, 27, 30, 49). It remains unclear what the reservoir of the fly- or diet-specific microorganisms is and how each is transferred between flies or to fresh diets, but our data make it clear that abundant fly microbiota ASVs are not necessarily sampled commonly from their diets. While there is ample evidence that the fly microbiota is assembled neutrally (6, 12, 50) and has priority effects (27), this suggests that either the flies are colonized early on with microorganisms that are not abundant in their diet; or that certain microorganisms are adept at invading established communities. Together, these findings are consistent with previous demonstrations that the host and community interactions drive the microbiota composition in the gut and the external fly environment (16, 29, 30, 45).

Along with host genotype and diet, abiotic environmental factors shape the *Drosophila* microbiota composition. We found that whereas no latitude was an inconsistent predictor of microbiota composition, the microbiota of flies sampled from different diets or years was individually correlated with environmental factors that influence microbiota composition. Of these, temperature was the strongest determinant. Elevated temperatures increased absolute and relative abundance of LAB in flies, consistent with their higher maximum growth temperatures than AAB. The tolerance of LAB to heat may also explain their elevated abundance in compost piles, which can reach internal temperatures above 60°C, relative to individual fallen fruits. However, high frequencies of LAB at warmer temperatures are inconsistent with our previously published expectation to detect LAB more abundantly in flies at high latitudes (5), but may possibly be explained by differences in the decomposition state of the diet. Also, a previous laboratory analysis of conventionally reared flies reported LAB were more abundant in flies reared at low temperatures (51). Therefore, one or more of our uses of gnotobiotic flies, which controls for the starting exposure to different microorganisms, or our use of different diets, host genotypes, or microorganisms may have contributed to the different trends. Regardless, temperature was a strong predictor of microbiota composition in the flies.

The various influences of temperature, photoperiod, humidity, diet type, and diet decomposition state, each of which can vary asynchronously with season in different geographic areas, raises questions about the role of seasonal variation in microbiota composition. Currently, the most comprehensive longitudinal analysis of the fruit fly microbiota comes from flies fed laboratory diets in outdoor mesocosms over a summer-to-fall season in the eastern USA (6). The flies’ microbiota clearly shifted with seasonal progression, including a summer-to-fall shift from *Acetobacter* to *Commensalibacter* dominance, and a mid-summer peak in the abundance of *Wautersiella*, a genus from the Flavobacteria that was not abundant in the flies we report on here (6). These seasonal patterns in the microbiota composition of flies reared outdoors were independent of variation in the diet since the flies were fed a laboratory diet throughout the experiment. Each of the samplings we report or reanalyze here were conducted over a relatively narrow ∼ 3 week window. We cannot speak to the other authors’ motivation, (**Fig. 1B, E**), but our intention in narrow sampling times was to control for time. With the retrospective perspective from the longitudinal analysis, our design actually increases variation by adding location-specific seasonal progression as an unreplicated confounding variable. It remains unclear exactly how seasonal progression should be defined, but our findings here suggest that temperature, time of fruit onset, and type of fruit available, plus humidity, day length, and UV irradiance as key factors to consider. If specific patterns in seasonal microbiota variation are common across multiple seasons, then such variation may also be a useful tool. For example, the relative abundances of *Acetobacter* and *Commensalibacter* in the **Fig. 1** datasets are sometimes positively, negatively, or not significantly correlated with each other (**Fig. S6**). One way of interpreting these varied outcomes is that the datasets where *Acetobacter* and *Commensalibacter* read counts are inversely related are asynchronous for seasonal progression; and datasets where these abundances are not significantly negatively correlated are more seasonally-synchronized. The strong influence of temperature alone suggests that timing a latitudinal sampling scheme based on predicted temperatures might be sufficient to account for this variation; although daily or site-specific deviations from the norm might it prohibitively difficult to synchronize. If the conditions are more varied, then such interpretations might require identifying additional seasonal trends that span years and locations. Regardless, the established patterns in seasonal evolution of *D. melanogaster* and the ability to track how variation in microbiota composition influences the fly suggest an ideal framework for future investigation of the relationship between seasonal variation, local adaptation of a host, and variation in microbiota composition.

The data presented here suggest the flies’ evolutionary history segregates with their diet and with their diet-dependent microbiota composition. We interpret these findings to mean that these factors together contribute phenotypic structure to wild populations even in the presence of expected gene flow. The distinct phenotypes of axenic flies from the different sampling locations confirms phenotypic differentiation between the flies collected from different diets at different geographic distances from each other, and that host genotype, as well as the microbiota, contributes to these differences. Beyond this, we observed wider differentiation of the development time trait between the flies at these sites when the microbiota was present. These latter findings are consistent with previous demonstrations that the microbiota can enhance phenotypic variance in wild fly populations (5), and likely underrepresent the potential variance because the flies were reared on the same diets and on a standardized microbiota composition.

This is because the constraints on diet and microbiota are both likely to be lower in the wild than in the laboratory. It is possible that the diet-dependent segregation of host phenotypes is mediated directly by host genotype based on variation in e.g., their dispersal, dietary or feeding preference traits (52–58). However, we note that there were no differences in host genetic selection on the microbiota in flies collected from compost or apples (**Fig. S8**), suggesting that the phenotypic differences observed in the wild flies are driven by diet or are sampling-specific, not from host genetic control of the microbiota (as in (21)). Further, the parallel phenotypic outcomes of flies reared in dispersal-limited mescosms with LAB, and wild in conditions that promote LAB abundance, are striking. When fly migration was restricted, flies feeding on *L. brevis* in their diets adopted a ‘faster’ life history strategy, weighing less, reaching larger population sizes, and developing to reproductive maturity more quickly than their *A. tropicalis-* fed counterparts (**Fig. 5C,** (1)). Flies feeding on compost in the wild, which promotes LAB abundance (**Fig. 2F**), mirrored these findings by displaying a shorter developmental period than flies collected from apples (**Fig. 5B**), a diet that restricts LAB prevalence and abundance (**Fig. 2F**). Together, the parallel outcomes of the two different lines of inquiry, one of which established the microbiota as a causal link, are consistent with the expectation that conditions that promote different fly microbiomes can shape their host’s evolutionary trajectory.

The analysis we present here does not focus on intracellular bacterium *Wolbachia*, a reproductive manipulator of many insects that is prevalent in wild *Drosophila* populations and can profoundly influence the flies’ life history and physiology (59–67). The major reason for this omission is that most of our datasets were poorly balanced for the presence and absence of *Wolbachia*; in some locations many flies bore *Wolbachia,* whereas *Wolbachia* were rare in other samplings. We generally discarded *Wolbachia* reads, then determined differences in microbiota composition between samples. However, we often reported *Wolbachia* colonization status (±) as a covariate in our analyses when it was significant, and performed unpublished comparisons of the *Wolbachia-*colonized and *Wolbachia*-free flies throughout this work. Most of the major statistical trends were observed when we analyzed the flies mixed together or separately according to their *Wolbachia* colonization status. Exceptions to this rule were generally confounded by small or unbalanced sample size and would require more sampling to make a conclusion either way, which is why we do not formally report on them here. As an exception, one dataset was well-balanced for *Wolbachia* colonization status: the analysis of flies collected on different fruits in a Middlefield, CT orchard (**TABLE S7**). Among these flies there were no genus-level differences in bacterial relative abundance between *Wolbachia* positive and - negative flies, but differences were detected at different taxonomic levels: *Wolbachia* positive flies bore more family-level *Acetobacteraceae* reads and, at the ASV level, more of a *Commensalibacter* ASV (**Fig. S7**). An association between *Wolbachia* and *Commensalibacter* was reported previously (6) but is somewhat surprising because a sister-genus, *Acetobacter*, has been reported by us and others to be negatively associated with *Wolbachia* abundance (5, 63, 68). These findings highlight the importance *Wolbachia* plays as a part of the *Drosophila* microbiota, and the recent use of outdoor enclosures to study *Wolbachia-Drosophila*-microbiota interactions highlights an elegant way to do so in a native setting while also maintaining a balanced statistical design.

In summary, we present evidence that many factors contribute to geographic variation in the *D. melanogaster* microbiota composition. We also show that identifying and considering such factors makes it clear that there are ordered patterns in the types and abundances of microorganisms in wild flies. In particular, several of our findings can help explain why LAB, which are among the most dominant taxa in flies reared on certain laboratory diets, are nearly absent from some recent surveys of flies in a wild setting. They also provide context supporting that members of the Enterobacteriaceae are common and abundant in flies in a wild setting.

Further analysis of these and other factors will improve our ability to explain and predict the many varied patterns of microbiota composition in these relatively simple communities of host-associated microorganisms, help to model the interactions in more complex communities of partners, and establish the role that microbial partners play in shaping animal life histories.

## MATERIALS AND METHODS

### General Rearing and Culture Conditions

We reared flies in general culture at 25°C on a 12-hour light:dark cycle. Flies from wild collections were reared on a molasses diet (5). Some laboratory experiments were performed using a CantonS fly line obtained from Mariana Wolfner that is *Wolbachia-*free and was reared on a yeast-glucose (Y-G) diet (23).

### Wild fly collections

We collected samples from eight orchards in the eastern USA in Fall of 2021 and five orchards in Utah, USA in Fall of 2020 (**Table S1**). At each location we collected samples from ten to twenty individual fallen fruit sites: the fruit, neighboring soil, a male *D. melanogaster* fly bearing a total microbiota and a male *D. melanogaster* fly bearing its resident microbiota. Flies were captured from an aerial insect net and divided into two empty, sterile vials; one immediately frozen on dry ice, the other left for > 2 hours for defecation of non-resident microorganisms and then frozen on dry ice. We aimed to collect 5-10 flies per vial at each fruit site. From each site, females were also collected and reared individually on molasses diet to establish isofemale lines that were kept in culture throughout the study. Additional fly populations were collected in 2023 following this method as well (**Table S2**). Collection and propagation of isofemale lines was as previously described, including morphological and molecular analysis to retain only *D. melanogaster* (45). Fruit and soil were sampled to ∼ 1/2-inch depth using a 1/4-inch diameter fruit corer that was briefly pre-sterilized in 10% bleach then pre-rinsed in sterile double-distilled H_2_O. Each were immediately stored on dry ice. We also performed aerial collections of flies at compost piles in three locations (**Table S1**), and single-fruit collections for the resident fly microbiota from many different apple, peach, and pear varieties in the orchard at Middlefield, CT. All samples were stored on dry ice until they could be permanently stored in a -80°C freezer.

Locale-specific fly populations were established from individual isofemale lines by 3 generations of common-garden mixing for 2-3 generations as described previously (45). Eggs laid by F2 or F3 flies were collected, dechorionated, and reared as gnotobiotic flies in association with the six bacterial species *Acetobacter tropicalis* DmCS_006*, Acetobacter sp.* DsW_54*, Acetobacter sp.* DmW_125*, Lactiplantibacillus plantarum* DmCS_001*, Weissella paramesenteroides* DmW_115, and *Leuconostoc suionicum* DmW_098. Two days after eclosion of adult flies, vials were transferred to new conditions to measure the influence of a 3-day perturbation in photoperiod (left at a 12h light:dark cycle or incubated at 1h light:23h dark or 23h light:1h dark) or temperature (left at 25°C or moved to 15 °C or 32 °C) on the adult fly microbiota. Three days later we measured the adult fly microbiota. Flies caught from compost and apples in the same orchard were reared under axenic and gnotobiotic conditions as individual isofemale lines, not in a common garden.

Experimentally-evolved flies were reared in outdoor mesocosms for 6.5 weeks as described previously (1), then flies from each mesocosm were reared in common garden populations for 2-3 generations. Bacteria-free embryos were derived from each population and their development time to adulthood was measured as described below.

### Gnobiotic fly rearing and enumeration of bacterial colonization

Embryos laid by *D. melanogaster* were collected, dechorionated in 10% bleach for two 150s washes, rinsed three times with sterile water, and 30-60 embryos were transferred to 7.5 ml sterile molasses diet in a 50 ml microcentrifuge tube. Separately, bacterial strains were cultured overnight in de Man-Rogosa-Sharpe (MRS) medium (Criterion C5932) for 1-2 days at 30°, normalized to OD_600_ = 0.1, and mixed in equal ratios. Then, we inoculated 50 μl of the bacterial mixture to the sterile eggs and allowed the eggs to develop to adulthood. When the adult flies were 4-7 days old, the adult fly microbiota was measured by collecting flies surface-sterilized in ethanol under light CO_2_ anesthesia, performing whole-body homogenization, and dilution plating the homogenate for CFU counting as described previously (27). From each fly vial we collected 4 pools of 2 male flies and 4 pools of 2 female flies. Fly homogenates were serially diluted onto MRS (to culture AAB and LAB) and MRS plus 10 μg/ml chloramphenicol and 10 μg/ml erythromycin (to culture only AAB). All dilution and plating was performed using an EpMotion 96. Plates were incubated at 30 °C for 2-3 days, and the antibiotic-free plates were cultures in sealed CO_2_-flooded containers. Colonies of LAB and AAB were then manually or automatically counted (27) to compare the abundances of each bacterial order in the fly microbiota. Each experiment was repeated three times with triplicate vials for each population and condition. CFU count data were analyzed as described previously (45). Significant influences of treatments on microbiota composition were determined by PERMANOVA of a Bray-Curtis beta-diversity distance matrix constructed from the LAB and AAB CFU counts, rarefied to 10000 (temperature) or 4000 (photoperiod) counts, using custom scripts, QIIME2, and the R package Vegan (69). PERMANOVA was always performed with 1,000 permutations. Variation in the absolute abundances of the LAB or AAB was tested for significance using a Kruskal-Wallis test with a post-hoc Dunn test and the R packages dunn.test (70), rcompanion (71), and multcomp (72).

### Sample preparation and 16S rRNA gene sequencing

In the laboratory, DNA was extracted from each sample using the Zymo Quick-DNA™ Fecal/Soil Microbe 96 Kit (D6011) following the manufacturer’s instructions except that DNA was eluted in 50 μl elution buffer. Flies were individually examined under a microscope to sequence only male *D. melanogaster*, as determined by the presence of a distinct genital arch. We also extracted DNA from 0.02 g of soil and 0.05 g of fruit. At the time DNA was extracted we loaded a microbiome cell standard onto a single well of each 96-well plate, and left three empty wells for a reagent-only control and two PCR amplification controls. We assured that negative PCR controls did not yield visible bands on gel electrophoreses.

We performed 16S rRNA marker gene sequencing using a previously described dual-barcoding approach (73). We normalized the reactions using the Just-a-Plate normalization kit (Charm Biotech, JN-120-10), and combined equivalent volumes of 96 samples into single pools. The pools were concentrated using a Zymo gDNA Clean & Concentrator 11-302C kit, fragment size distributions were evaluated on an Agilent FemtoPulse (Agilent Technologies, Santa Clara, CA, USA), and fragments in the 250-450 bp range were selected on a Sage Science Blue Pippin (Sage Science, Beverly, MA, USA). The final molarity of the pool was estimated via qPCR at the BYU DNA sequencing center. Then, we combined the pools and sequenced them on an Illumina MiSeq using 500 cycle v2 chemistry as described previously (73).

### Sequence analysis

Demultiplexed sequence reads were analyzed using QIIME2 (74) and R 4.1.2 (75). Previously published datasets were accessed from our own personal archives (**Fig. 1B** (5)), though the data are published at PRJNA589709, or from BioProject <u>PRJNA873107</u>. Using DADA2 (76), reads were denoised, dereplicated, and amplicon sequence variants (ASVs) were called using trimming lengths that maximize quality scores of the reads. Taxonomic assignments to ASV were made using the GreenGenes classifier 13_8_99 (77), and samples were assigned as *Wolbachia* positive if 20% or more of the total reads were assigned to *Wolbachia.* Then, reads that could be assigned to *Archaea, Chloroplast, Mitochondria,* or *Wolbachia* were discarded. Operational taxonomic unit (OTU) tables were filtered to various thresholds per sample, reported in each corresponding figure legend. Significant differences between groups were determined by PERMANOVA (78) of beta-diversity distance metrics (79, 80). Phylogenetic trees, supporting the use of the Unifrac distance metrics, were built with fasttree2 (81) based on mafft alignment (82). Analysis of Composition of Microbiomes (ANCOM) was used to define significant differences in the abundances of individual microbes between samples (83). ANCOM was performed on reads clustered at each taxonomic level and is generally reported at the genus or ASV level only, though trends across levels were usually consistent. Beta-diversity distance analysis, calculation of Venn diagrams (84), Spearman’s rank correlation tests, and PERMANOVA (78) were all performed in R. Application of neutral models to sequencing data were performed in R using default parameters as described previously (85, 86), and Akaike information criterion (AIC) values for each test were recorded from each separate model. To perform Mantel tests, in R, environmental metadata for each location were downloaded from the USA & Americas (1998-2022) database at https://nsrdb.nrel.gov/data-viewer in 30- or 60-minute intervals or from the Meteostate Prime Meridian: Africa and Europe database in 60 minute intervals. Maximum, average, or minimum values of each character were calculated on the day of sampling, a distance matrix was constructed based on individual or multiple values, and the relationship between the sampling location of each individual sample and the environmental metadata were calculated using a Mantel test based on Spearman’s rank correlations. Sequences from the DEST dataset were downloaded from the SRA using fasterq-dump v.2.10.8 from the SRA toolkit (https://github.com/ncbi/sra-tools), and microbial profiling was performed using MetaPhlan3 and the ChocoPhlan v30 database (87). The OTU table obtained from all outputs was then analyzed using Bray Curtis distance metrics as described above.

### Rearing gnotobiotic flies on fruit diets

To measure the impact of distinct diets on the microbiota composition of *D. melanogaster*, we reared gnotobiotic *D. melanogaster* CantonS on ten types of fruit, obtained from grocery stores in Provo and Orem, UT. We diced, froze, and transferred 5-10 g of each fruit type to a 50 ml centrifuge tube, then autoclaved the tubes. Then, we transferred 30-60 bleach-sterilized, dechorionated fly embyos to the diets as described above, and inoculated each vial with a mixture of separately cultured and OD_600_=0.1-normalized bacterial community composed of *Acetobacter pomorum* DmCS_004, *Acetobacter tropicalis* DmCS_006, *Bacillus subtilis* 168, *Escherichia coli* K-12 MG1655, *Gluconobacter cerinus* Dm-58, *Komagataeibacter medellinensis* NBRC 3288*, Lactiplantibacillus plantarum* DmCS_001, *Levilactobacillus brevis*, *Pantoea* sp. JGM49, *Pantoea dispersa* JGM106, DmCS_003, *Providencia rettgeri* JGM232, *Weissella paramesenteroides* DmW_115. When the flies were 5-7 day old adults, one pool of 5 male flies was collected from each vial, frozen at -80 °C, and the V4 region of the 16S rRNA gene was sequenced and analyzed as described above. We performed four separate experiments, each with triplicate vials of flies, to target collection of 3 pools of flies in each of three experiments. However, recovery of adult flies was often challenging because some diets poorly supported fly growth or were wet and adult flies drowned. We collected and sequenced as many samples as we could recover from the four experiments.

### Rearing wild flies on decomposing diets

To dissect the separate influences of seasonal progression and diet decomposition on microbiota composition, we established fresh fruit piles in a field site in Provo, UT in August 2022. Each pile was established by dropping one bushel of fruit (Allred’s Orchards, Payson, UT), from a height of 4 feet onto a grass lawn. We established piles from apples and peaches every two weeks for six weeks, and a fourth apple pile two weeks later (fresh peaches were no longer available). Twice a week up to 30 fruit flies were sampled from each pile with a hand insect vacuum into empty vials, starved for 2 hours, then frozen in a -80 °C freezer. Sampling continued until hard freeze onset, November 2022, and the 16S rRNA V4 region on individual flies was sequenced as described above. Only 7 of 146 male flies were assigned as *D. simulans* by morphological examination, so we analyzed male *D. melanogaster* and female *Drosophila* together in this experiment.

### Measuring fly development time to adulthood

Fly development time to reproductive maturity was measured at 1, 6, and 11 hours into the daily light cycle each day as the time to eclosion for each individual pupa on the side of a fly vial until all flies in a vial had eclosed or there were two consecutive time points where no flies eclosed, whichever came first. Significant differences in fly development time were determined using a Cox mixed-effects survival model with the source diet (apples or compost) and rearing treatment (axenic or gnotobiotic) as interative terms and experimental replicate (four fly vials per treatment were reared in each of three distinct experiments in time) as a random effect. The analyses were performed using survival-model specific R packages (88, 89).

### Data analysis

Some R packages we used are not cited elsewhere (90–103). Maps were made in R using (104). A screenshot from Apple Maps taken in summer 2024 was used in **Fig. 5A**. Raw data and scripts for these analysis are available at https://github.com/johnchaston/Gale2024, most prominently the script file Gale_script.Rmd and the knitted HTML file from that script, Gale_script.html.zip.

### Data availability

Sequences from this study will be deposited and made publically available in the SRA at accession XXXXX when the article is accepted.

## ACKNOWLEDGEMENTS

Research reported in this publication was supported in part by the National Institute of General Medical Sciences of the National Institutes of Health under Award Number R15GM140388. The content is solely the responsibility of the authors and does not necessarily represent the official views of the National Institutes of Health.

