## Supplementary material for "Environment and diet shape the geography-specific *Drosophila melanogaster* microbiota composition": SI appendix

### Table S1. Sampling site details

| CITY | ORCHARD NAME | LATITUDE | LONGITUDE | SEPARATE COMPOST COLLECTION | # LINES | DATE COLLECTED | FRUIT TYPE |
|---|---|---|---|---|---|---|---|
| **CHARLOTTESVILLE, VA** | Carter Mountain Orchard | 38.07474 | -78.38155 | | 17 | 10/20/21 | Apple |
| **CHURCHVILLE, MD** | Lohr's Orchard | 39.539 | -76.2399 | | 12 | 10/21/21 | Apple |
| **MEDIA, PA** | Indian Orchards, Linvilla Orchards | 39.8845 | -75.4125 | | 28 | 10/28/21 | Apple |
| **MIDDLEFIELD, CT** | Lyman Orchards | 41.49383 | -72.71145 | Y | 13 | 10/1/21 | Apple; separately, peaches and pears |
| **HARVARD, MA** | Westward Orchards | 42.50867 | -71.57071 | Y | 10 | 9/30/21 | Apple |
| **DURHAM, NH** | Demeritt Hill Farm | 43.08745 | -71.04666 | | 24 | 9/30/21 | Apple |
| **BOWDOIN, ME** | Rocky Ridge Orchard | 44.0253 | -69.94379 | | 19 | 9/29/21 | Apple |
| **ETNA, ME** | Conant Orchards | 44.82082 | -69.11144 | Y | ~20 | 9/29/21 | Apple |
| **SANTAQUIN, UT** | Rowley's Red Barn | 39.966865 | -111.791562 | | na | 10/10/20 | Peach |
| **LINDON, UT** | Martel Famrs | 40.3327707 | -111.7219739 | | na | 9/29/20, 10/6/20, 10/10/20 | Peach |
| **ALPINE, UT** | Burgess Orchards | 40.4457589 | -111.7818936 | | na | 10/9/20 | Peach |
| **OGDEN, UT** | Stake Farm North Ogden Peach Orchard | 41.326252 | -112.011375 | | na | 10/9/20 | Peach |
| **LOGAN, UT** | Zollinger Fruit and Tree Farm | 41.72573 | --111.809883 | | na | 10/8/20 | Peach |

**Table S2 Sampling site details for individual isofemale lines**

| LINE NAME | ORCHARD | LATITUDE | LONGITUDE | DATE COLLECTED | FRUIT TYPE |
|---|---|---|---|---|---|
| **TW** | Conant Orchards | 44.82082 | -69.11144 | 10-26-2023 | Apple Δ[a] |
| **TI** | Conant Orchards | 44.82082 | -69.11144 | 10-26-2023 | Apple Δ |
| **ER** | Conant Orchards | 44.82082 | -69.11144 | 10-26-2023 | Apple ❍ [a] |
| **P19** | Conant Orchards | 44.82082 | -69.11144 | 9-30-2023 | Compost ☐ [a] |
| **P20** | Conant Orchards | 44.82082 | -69.11144 | 9-30-2023 | Compost ☐ |
| **P21** | Conant Orchards | 44.82082 | -69.11144 | 10-26-2023 | Compost ☐ |

[a] symbol position shown in Figure 5A

**Table S3. Significant covariance by a Mantel test between a single value and Bray-Curtis distance in sequencing datasets. Cells contain Mantel test p-value. Bold represents p < 0.05.**

| | *Sampling experiment* | | | | | | |
|---|---|---|---|---|---|---|---|
| ***Environmental parameter*** | 2009 | 2018 Vineyard | 2018 Orchard | 2021 | 2020 Utah | DEST N. A. | DEST Europe |
| *Daily maximum temperature* | **0.003** | **0.002** | **0.041** | **0.001** | **0.007** | **0.019** | **0.004** |
| *Daily minimum relative humidity* | **0.003** | **0.001** | **0.021** | **0.001** | 0.065 | 0.29 | **0.012** |
| *Minimum precipitable water* | **0.001** | **0.002** | **0.022** | **0.002** | 0.166 | 0.14 | 0.35 |
| *Daily maximum Clearsky direct normal irradiance* | **0.003** | **0.001** | **0.024** | **0.033** | 0.125 | 0.07 | 0.12 |
| *Daily average wind direction* | **0.045** | **0.002** | **0.033** | **0.001** | 0.089 | 0.60 | 0.24 |
| *Daily average relative humidity* | **0.001** | **0.001** | 0.053 | **0.001** | 0.223 | 0.20 | **0.013** |
| *Daily maximum Clearsky global horizontal irradiance* | **0.029** | **0.041** | 0.05 | **0.011** | 0.312 | **0.016** | **0.002** |
| *Daily minimum temperature* | **0.001** | **0.015** | 0.095 | **0.001** | 0.376 | **0.038** | 0.055 |
| *Daily maximum dew point* | **0.001** | **0.003** | 0.155 | **0.045** | 0.39 | 0.26 | 0.15 |
| *Daily average dew point* | **0.015** | **0.001** | 0.168 | **0.002** | 0.11 | 0.24 | 0.26 |
| *Daily average diffuse horizontal irradiance* | **0.001** | **0.017** | 0.182 | **0.001** | 0.471 | 0.94 | 0.21 |
| *Daily maximum diffuse horizontal irradiance* | **0.011** | **0.001** | 0.221 | **0.006** | 0.312 | 0.97 | 0.35 |
| *Daily maximum wind speed* | **0.001** | **0.001** | 0.225 | **0.008** | **0.012** | 0.061 | 0.11 |
| *Daily average windspeed* | **0.002** | **0.001** | 0.248 | **0.045** | **0.029** | 0.087 | 0.23 |
| *Daily minimum wind speed* | **0.001** | **0.001** | 0.354 | 0.054 | 0.19 | 0.18 | 0.39 |
| *Daily average temperature* | **0.001** | 0.059 | 0.071 | **0.001** | 0.591 | **0.03** | **0.003** |
| *Daily minimum wind direction* | **0.001** | 0.087 | 0.257 | **0.014** | **0.029** | 0.78 | 0.57 |
| *Daily minimum dew point* | **0.001** | 0.137 | 0.086 | **0.001** | 0.242 | 0.24 | 0.36 |
| *Daily average precipitable water* | **0.001** | 0.202 | **0.013** | **0.005** | 0.277 | 0.18 | 0.097 |
| *Daily average Clearsky direct normal irradiance* | **0.015** | 0.276 | 0.261 | **0.016** | 0.316 | 0.16 | **0.005** |
| *Daily maximum precipitable water* | **0.035** | 0.324 | **0.03** | **0.001** | 0.261 | 0.15 | **0.037** |
| *Daily maximum relative humidity* | **0.021** | NA | 0.064 | **0.001** | 0.513 | 0.27 | 0.74 |
| *Daily maximum Clearsky diffuse horizontal irradiance* | 0.058 | 0.07 | 0.206 | **0.001** | 0.069 | 0.69 | 0.62 |
| *Daily average Clearsky global horizontal irradiance* | 0.06 | **0.028** | 0.219 | **0.005** | 0.357 | **0.046** | **0.001** |
| *Daily average Clearsky diffuse horizontal irradiance* | 0.083 | 0.178 | 0.191 | **0.001** | 0.085 | 0.77 | 0.19 |
| *Daily average surface albedo* | 0.15 | 0.987 | **0.043** | **0.002** | **0.013** | 0.37 | 0.76 |
| *Daily maximum surface albedo* | 0.15 | 0.987 | **0.043** | **0.002** | **0.013** | 0.45 | 0.75 |
| *Daily minimum surface albedo* | 0.15 | 0.987 | **0.043** | **0.002** | **0.013** | 0.48 | 0.75 |
| *Daily average pressure* | 0.18 | 0.957 | **0.045** | 0.126 | **0.008** | **0.026** | 0.094 |
| *Daily maximum fill flag* | 0.247 | **0.002** | 0.279 | **0.001** | 0.143 | 0.53 | 0.79 |
| *Daily maximum pressure* | 0.248 | 0.879 | 0.133 | 0.114 | **0.008** | **0.031** | 0.08 |
| *Daily minimum pressue* | 0.303 | 0.94 | **0.044** | 0.131 | **0.016** | **0.022** | 0.09 |
| *Dailiy average solar zenith angle* | 0.491 | **0.027** | 0.359 | **0.001** | 0.437 | **0.009** | **0.001** |
| *Daily maximum solar zenith angle* | 0.577 | 0.531 | 0.168 | **0.001** | 0.329 | **0.009** | **0.019** |
| *Daily average fill flag* | 0.615 | 0.95 | 0.285 | **0.002** | 0.088 | 0.072 | 0.11 |
| *Daily minimum solar zenith angle* | 0.71 | **0.003** | 0.378 | **0.001** | 0.226 | **0.019** | **0.001** |
| *Daily maximum wind direction* | 0.988 | 0.566 | **0.017** | **0.001** | **0.011** | 0.16 | 0.20 |

**Table S4 PERMANOVA results for CFU counts (AAB and LAB) of the fly microbiota (see Fig 1N, 1P)**

| | Absolute | | | | | | Relative | | | | | |
|---|---|---|---|---|---|---|---|---|---|---|---|---|
| | Df[a] | SS[b] | $R^2$ | F | p[c] | | Df[a] | SS[b] | $R^2$ | F | p[c] | |
| Geography | 7 | 20.94 | 0.10 | 19.01 | 0.001 | *** | 7 | 8.50 | 0.14 | 29.75 | 0.001 | *** |
| Temperature | 2 | 15.45 | 0.07 | 49.10 | 0.001 | *** | 2 | 4.81 | 0.08 | 58.90 | 0.001 | *** |
| Sex | 1 | 0.46 | 0.00 | 2.93 | 0.03 | * | 1 | 0.12 | 0.00 | 2.99 | 0.07 | . |
| Exp[d] | 2 | 9.93 | 0.05 | 31.54 | 0.001 | *** | 2 | 1.57 | 0.03 | 19.21 | 0.001 | *** |
| G * T[e] | 13 | 6.36 | 0.03 | 3.11 | 0.001 | *** | 13 | 1.56 | 0.03 | 2.93 | 0.001 | *** |
| G * S | 7 | 1.00 | 0.00 | 0.91 | 0.60 | | 7 | 0.39 | 0.01 | 1.36 | 0.20 | |
| T * S | 2 | 1.39 | 0.01 | 4.42 | 0.002 | ** | 2 | 0.19 | 0.00 | 2.28 | 0.10 | |
| E / Plate[f] | 3 | 2.84 | 0.01 | 6.01 | 0.001 | *** | 3 | 0.29 | 0.00 | 2.35 | 0.07 | . |
| G * T * S | 13 | 2.05 | 0.01 | 1.00 | 0.48 | | 13 | 0.85 | 0.01 | 1.60 | 0.07 | . |
| E / P / Vial[g] | 81 | 35.37 | 0.16 | 2.78 | 0.001 | *** | 81 | 10.98 | 0.19 | 3.32 | 0.001 | *** |
| Residual | 762 | 119.92 | 0.56 | | | | 738 | 30.10 | 0.51 | | | |
| Total | 893 | 215.71 | 1.00 | | | | 869 | 59.33 | 1.00 | | | |
| Geography | 7 | 13.11 | 0.05 | 14.45 | 0.001 | *** | 7 | 1.76 | 0.03 | 7.42 | 0.001 | *** |
| Photoperiod | 2 | 1.50 | 0.01 | 5.77 | 0.001 | *** | 2 | 0.68 | 0.01 | 9.99 | 0.001 | *** |
| Sex | 1 | 0.20 | 0.00 | 1.53 | 0.181 | | 1 | 0.04 | 0.00 | 1.28 | 0.253 | |
| Exp[d] | 2 | 17.35 | 0.07 | 66.90 | 0.001 | *** | 2 | 10.69 | 0.16 | 157.70 | 0.001 | *** |
| G * P[e] | 14 | 6.16 | 0.02 | 3.39 | 0.001 | *** | 14 | 1.50 | 0.02 | 3.15 | 0.001 | *** |
| G * S | 7 | 1.15 | 0.00 | 1.26 | 0.187 | | 7 | 0.60 | 0.01 | 2.54 | 0.013 | * |
| P * S | 2 | 0.57 | 0.00 | 2.21 | 0.033 | * | 2 | 0.03 | 0.00 | 0.45 | 0.639 | |
| E / Plate[f] | 3 | 5.07 | 0.02 | 13.03 | 0.001 | *** | 3 | 2.58 | 0.04 | 25.36 | 0.001 | *** |
| G * P * S | 14 | 2.29 | 0.01 | 1.26 | 0.119 | | 14 | 0.74 | 0.01 | 1.55 | 0.086 | . |
| E / Plate / Vial[g] | 141 | 59.31 | 0.23 | 3.24 | 0.001 | *** | 141 | 10.31 | 0.15 | 2.16 | 0.001 | *** |
| Residual | 1207 | 156.50 | 0.59 | | | | 1144 | 38.78 | 0.57 | | | |
| Total | 1400 | 263.20 | 1.00 | | | | 1337 | 67.71 | 1.00 | | | |

[a] degrees of freedom

[b] sum of squares

[c] p-value

[d] One of three separate experiments in time

[e] “*” = the interaction term

[f] “/” = the nesting term; Plate = the 96-well plate on which samples were homogenized from which they were dilution plated

[g] vial = the source vial the flies were grown in (30-50 mixed sex flies per vial)

### Table S5 PERMANOVA results for 16S sequencing data corresponding to Figure 2A-B

| | *Unweighted Unifrac* | | | | | | *Weighted Unifrac* | | | | | *Bray Curtis* | | | | |
|---|---|---|---|---|---|---|---|---|---|---|---|---|---|---|---|---|
| | **Df**[a] | **SS**[b] | **$R^2$** | **F** | **p**[c] | | **SS** | **$R^2$** | **F** | **p** | | **SS** | **$R^2$** | **F** | **p** | |
| *Sample type*[d] | 3 | 19.01 | 0.37 | 45.97 | 0 | *** | 8.65 | 0.36 | 44.33 | 0 | *** | 17.9 | 0.2 | 20.08 | 0 | *** |
| *Orchard* | 7 | 2.08 | 0.04 | 2.16 | 0 | *** | 0.87 | 0.04 | 1.9 | 0.02 | * | 4.74 | 0.05 | 2.28 | 0 | *** |
| *Individual fruit* | 66 | 10.03 | 0.19 | 1.1 | 0.15 | | 4.48 | 0.19 | 1.04 | 0.35 | | 21.45 | 0.24 | 1.09 | 0.04 | * |
| *S * O* [e] | 18 | 3.71 | 0.07 | 1.5 | 0 | *** | 1.73 | 0.07 | 1.48 | 0.04 | * | 9.91 | 0.11 | 1.85 | 0 | *** |
| *Residual* | 124 | 17.09 | 0.33 | NA | NA | | 8.06 | 0.34 | NA | NA | | 36.84 | 0.41 | NA | NA | |
| *Total* | 218 | 51.92 | 1 | NA | NA | | 23.79 | 1 | NA | NA | | 90.85 | 1 | NA | NA | |

[a] degrees of freedom

[b] sum of squares

[c] p-value

[d] Flies (total), flies (resident), fruit, or soil

[e] “*” = the interaction term

### Table S6 PERMANOVA results for 16S sequencing data corresponding to Figure 2F

| | Unweighted Unifrac | | | | | | Weighted Unifrac | | | | | Bray Curtis | | | | |
|---|---|---|---|---|---|---|---|---|---|---|---|---|---|---|---|---|
| | Df[a] | SS[b] | $R^2$ | F | p[c] | | SS | $R^2$ | F | p | | SS | $R^2$ | F | p | |
| *Fresh or compost* | 1 | 0.56 | 0.1 | 4.49 | $<10^{-4}$ | *** | 0.11 | 0.08 | 3.53 | 0.02 | * | 0.87 | 0.07 | 3.25 | $<10^{-4}$ | *** |
| *Orchard* | 2 | 0.44 | 0.08 | 1.78 | 0.03 | * | 0.12 | 0.08 | 1.89 | 0.1 | | 1.71 | 0.14 | 3.2 | $<10^{-4}$ | *** |
| *F * O* [d] | 2 | 0.28 | 0.05 | 1.13 | 0.29 | | 0.17 | 0.12 | 2.77 | 0.03 | * | 1.26 | 0.1 | 2.35 | $<10^{-4}$ | *** |
| *Residual* | 33 | 4.1 | 0.76 | NA | NA | | 1.02 | 0.72 | NA | NA | | 8.82 | 0.7 | NA | NA | |
| *Total* | 38 | 5.38 | 1 | NA | NA | | 1.41 | 1 | NA | NA | | 12.65 | 1 | NA | NA | |

[a] degrees of freedom

[b] sum of squares

[c] p-value

[d] "*" = the interaction term

### Table S7 PERMANOVA results for 16S sequencing data corresponding to Figure 3A

| | *Unweighted Unifrac* | | | | | | *Weighted Unifrac* | | | | | *Bray Curtis* | | | | |
|---|---|---|---|---|---|---|---|---|---|---|---|---|---|---|---|---|
| | **Df**[a] | **SS**[b] | **$R^2$** | **F** | **p**[c] | | **SS** | **$R^2$** | **F** | **p** | | **SS** | **$R^2$** | **F** | **p** | |
| *Fruit type* [d] | 2 | 0.43 | 0.08 | 1.36 | 0.13 | | 0.12 | 0.10 | 2.41 | 0.04 | * | 1.08 | 0.10 | 2.04 | $< 10^{-4}$ | *** |
| *Wolbachia* | 1 | 0.32 | 0.06 | 2.00 | 0.03 | * | 0.14 | 0.11 | 5.32 | $< 10^{-4}$ | *** | 0.56 | 0.05 | 2.11 | 0.01 | * |
| *F / Variety* [e] | 7 | 1.34 | 0.25 | 1.20 | 0.15 | | 0.37 | 0.29 | 2.07 | 0.02 | * | 2.69 | 0.26 | 1.45 | 0.01 | * |
| *F * W* [f] | 2 | 0.21 | 0.04 | 0.68 | 0.9 | | 0.08 | 0.06 | 1.59 | 0.16 | | 0.86 | 0.08 | 1.62 | 0.01 | * |
| *( F / V ) * W* | 5 | 0.75 | 0.14 | 0.94 | 0.58 | | 0.21 | 0.17 | 1.67 | 0.09 | | 1.55 | 0.15 | 1.17 | 0.19 | |
| *Residual* | 14 | 2.23 | 0.42 | NA | NA | | 0.36 | 0.28 | NA | NA | | 3.71 | 0.36 | NA | NA | |
| *Total* | 31 | 5.28 | 1.00 | NA | NA | | 1.29 | 1.00 | NA | NA | | 10.46 | 1.00 | NA | NA | |

[a] degrees of freedom
[b] sum of squares
[c] p-value
[d] apples, peaches, or pears
[e] "/" = the nesting term
[f] "*" = the interaction term

### Table S8 PERMANOVA results for 16S sequencing data corresponding to Figure 3B

| | *Unweighted Unifrac* | | | | | *Weighted Unifrac* | | | | | *Bray Curtis* | | | | |
|---|---|---|---|---|---|---|---|---|---|---|---|---|---|---|---|
| | **Df**[a] | **SS**[b] | **$R^2$** | **F** | **p**[c] | **SS** | **$R^2$** | **F** | **p** | | **SS** | **$R^2$** | **F** | **p** | |
| *Fruit* | 9 | 0.6 | 0.21 | 1.24 | 0.08 | 0.53 | 0.34 | 2.42 | $<10^{-4}$ | *** | 5.16 | 0.36 | 2.67 | $<10^{-4}$ | *** |
| *Residual* | 43 | 2.3 | 0.79 | NA | NA | 1.05 | 0.66 | NA | NA | | 9.24 | 0.64 | NA | NA | |
| *Total* | 52 | 2.89 | 1 | NA | NA | 1.58 | 1 | NA | NA | | 14.4 | 1 | NA | NA | |

[a] degrees of freedom

[b] sum of squares

[c] p-value

### Table S9 PERMANOVA results for 16S sequencing data corresponding to Figure 4A, Figure 4H

| | *Unweighted Unifrac* | | | | | | *Weighted Unifrac* | | | | | *Bray Curtis* | | | | |
|---|---|---|---|---|---|---|---|---|---|---|---|---|---|---|---|---|
| | **Df**[a] | **SS**[b] | **$R^2$** | **F** | **p**[c] | | **SS** | **$R^2$** | **F** | **p** | | **SS** | **$R^2$** | **F** | **p** | |
| *Fruit type* | 1 | 0.96 | 0.03 | 6.64 | $<10^{-4}$ | *** | 0.18 | 0.04 | 8.72 | $<10^{-4}$ | *** | 3.21 | 0.05 | 11.79 | $<10^{-4}$ | *** |
| *Calendar date* | 1 | 0.78 | 0.03 | 5.41 | $<10^{-4}$ | *** | 0.22 | 0.05 | 10.5 | $<10^{-4}$ | *** | 3.03 | 0.05 | 11.15 | $<10^{-4}$ | *** |
| *Fly sex* | 1 | 0.43 | 0.01 | 3 | $<10^{-4}$ | *** | 0.04 | 0.01 | 1.68 | 0.12 | | 0.69 | 0.01 | 2.53 | $<10^{-4}$ | *** |
| *Wolbachia* | 1 | 0.26 | 0.01 | 1.8 | 0.06 | | 0.05 | 0.01 | 2.36 | 0.05 | | 0.49 | 0.01 | 1.81 | 0.03 | |
| *F / pile* [d] | 5 | 1.49 | 0.05 | 2.07 | $<10^{-4}$ | *** | 0.14 | 0.03 | 1.36 | 0.12 | | 5.4 | 0.09 | 3.97 | $<10^{-4}$ | *** |
| *F * C* [e] | 1 | 0.5 | 0.02 | 3.45 | $<10^{-4}$ | *** | 0.02 | 0 | 0.74 | 0.56 | | 0.97 | 0.02 | 3.57 | $<10^{-4}$ | *** |
| *( F / P ) * C* | 5 | 1.08 | 0.04 | 1.5 | 0.02 | * | 0.2 | 0.05 | 1.94 | 0.04 | * | 3.2 | 0.05 | 2.36 | $<10^{-4}$ | *** |
| *Residuals* | 168 | 24.17 | 0.81 | NA | NA | | 3.53 | 0.81 | NA | NA | | 45.7 | 0.73 | NA | NA | |
| *Total* | 183 | 29.67 | 1 | NA | NA | | 4.38 | 1 | NA | NA | | 62.7 | 1 | NA | NA | |
| *Fruit type* | 1 | 0.96 | 0.03 | 6.64 | $<10^{-4}$ | *** | 0.18 | 0.04 | 8.72 | $<10^{-4}$ | *** | 3.21 | 0.05 | 11.79 | $<10^{-4}$ | *** |
| *Establishment time* | 1 | 0.75 | 0.03 | 5.21 | $<10^{-4}$ | *** | 0.09 | 0.02 | 4.06 | 0.01 | * | 2.04 | 0.03 | 7.48 | $<10^{-4}$ | *** |
| *Sex* | 1 | 0.42 | 0.01 | 2.92 | $<10^{-4}$ | *** | 0.03 | 0.01 | 1.63 | 0.14 | * | 0.7 | 0.01 | 2.59 | $<10^{-4}$ | *** |
| *Wolbachia* | 1 | 0.22 | 0.01 | 1.54 | 0.11 | | 0.04 | 0.01 | 2.06 | 0.07 | | 0.42 | 0.01 | 1.54 | 0.08 | |
| *F / pile* | 5 | 1.57 | 0.05 | 2.18 | $<10^{-4}$ | *** | 0.29 | 0.07 | 2.72 | $<10^{-4}$ | *** | 6.46 | 0.1 | 4.75 | $<10^{-4}$ | *** |
| *F * E* | 1 | 0.5 | 0.02 | 3.45 | $<10^{-4}$ | *** | 0.02 | 0 | 0.74 | 0.56 | | 0.97 | 0.02 | 3.57 | $<10^{-4}$ | *** |
| *( F / P ) * E* | 5 | 1.08 | 0.04 | 1.5 | 0.02 | * | 0.2 | 0.05 | 1.94 | 0.04 | * | 3.2 | 0.05 | 2.36 | $<10^{-4}$ | *** |
| *Residuals* | 168 | 24.17 | 0.81 | NA | NA | | 3.53 | 0.81 | NA | NA | | 45.7 | 0.73 | NA | NA | |
| *Total* | 183 | 29.67 | 1 | NA | NA | | 4.38 | 1 | NA | NA | | 62.7 | 1 | NA | NA | |

[a] degrees of freedom

[b] sum of squares

[c] p-value

[d] "/" = the nesting term; Plate = the 96-well plate on which samples were homogenized from which they were dilution plated

[e] "*" = the interaction term

### Table S10 PERMANOVA results for CFU data corresponding to Figure S8

| | Unrarefied Bray Curtis (Absolute abundance) | | | | | | Rarefied Bray Curtis (Relative abundance) | | | | |
|---|---|---|---|---|---|---|---|---|---|---|---|
| | Df[a] | SS[b] | $R^2$ | F | p[c] | | SS | $R^2$ | F | p | |
| *Source (apples or compost)* | 1 | 0.31 | 0.01 | 1.54 | 0.16 | | 0.00 | 0.00 | 0.08 | 0.78 | |
| *Fly sex* | 1 | 1.20 | 0.05 | 5.98 | 0.001 | *** | 0.19 | 0.03 | 3.72 | 0.06 | . |
| *Experimental replicate* | 1 | 3.71 | 0.17 | 18.44 | 0.001 | *** | 2.37 | 0.38 | 47.52 | 0.001 | *** |
| *So * Se* [d] | 1 | 0.15 | 0.01 | 0.77 | 0.57 | | 0.10 | 0.02 | 2.04 | 0.16 | |
| *Residual* | 82 | 16.50 | 0.75 | | | | 3.59 | 0.57 | | | |
| *Total* | 86 | 21.87 | 1.00 | | | | 6.26 | 1.00 | | | |

[a] degrees of freedom

[b] sum of squares

[c] p-value

[d] "*" = the interaction term

**Figure S1. The relationship between microbiota composition and latitude in flies sampled from Utah in Fall 2020.** A) Sampling map. B) Taxon plot. Plots showing the correlation between Bray Curtis distance (a measure of beta-diversity in microbiota composition) and environmental distance defined by C) latitude, and D) temperature. Mantel test results rho ( ρ) and p-value (p) are also shown. Read counts were rarefied to 500 reads prior to analysis.

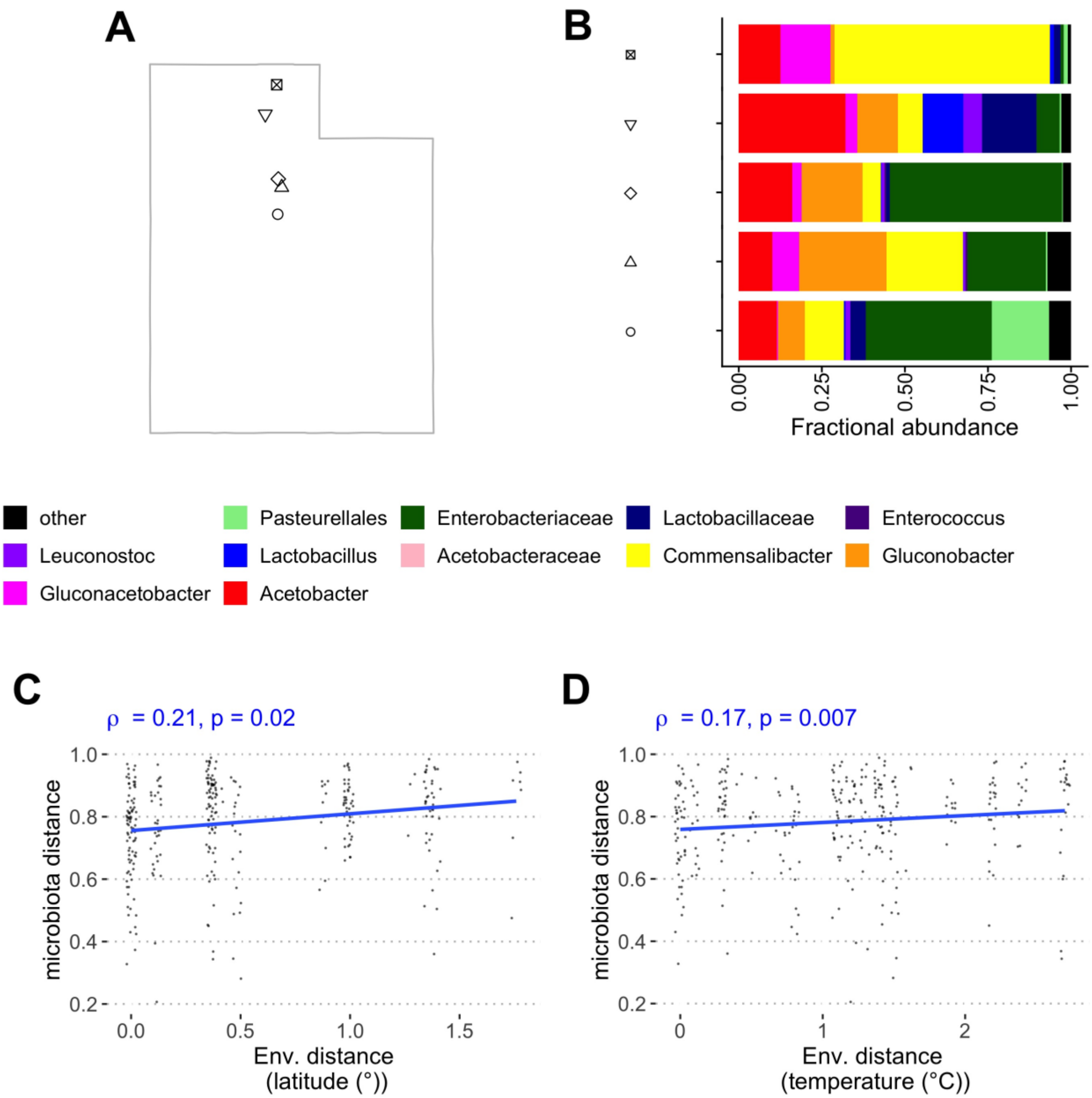

**Figure S2. The correlation between microbiota composition and environmental distance.** *Data correspond to Figure 1B-M*. Samples were collected from the eastern USA in A,E,I,M) 2009 (from apples and peaches), B,F,J,N) 2018 (from grapes), C,G,K,O) 2018 (from apples), D, H, L, P) 2021 (from apples). Environmental distance was defined as shown on each x-axis. Microbiota distance was defined using the A-H) weighted or I-P) unweighted Unifrac distance metric.

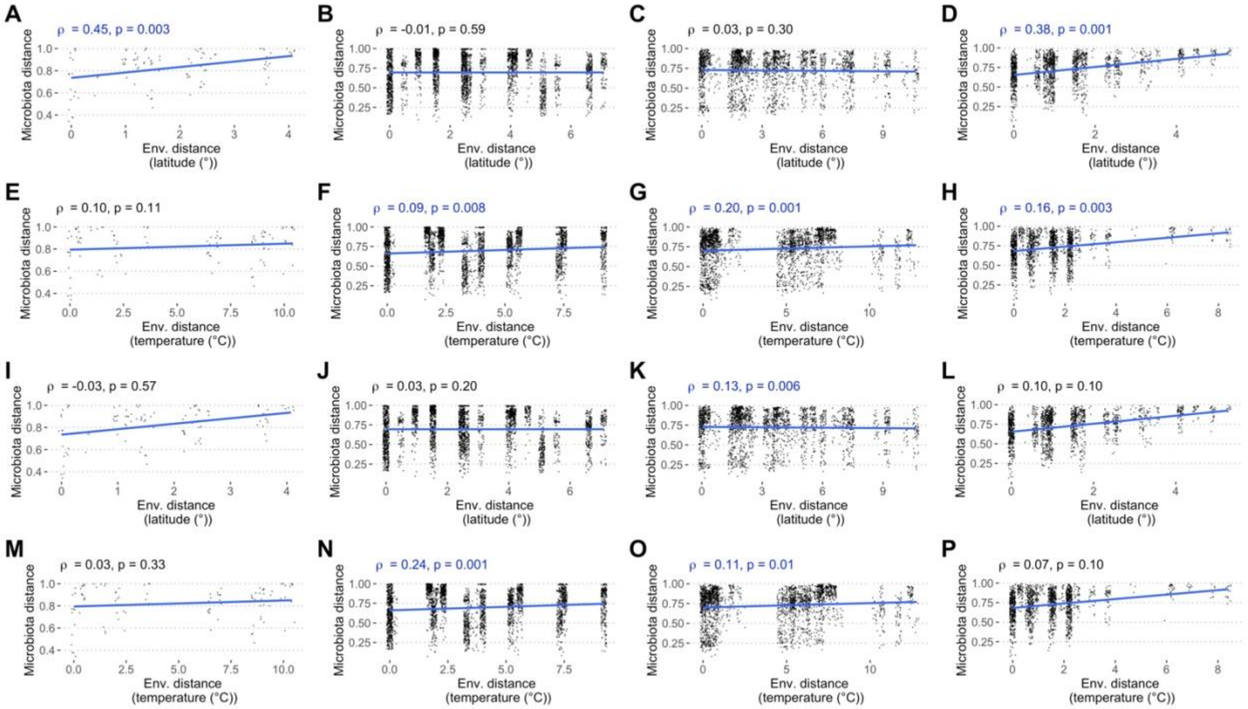

**Figure S3. Temperature-dependent variation in the microbiota composition of common garden fly populations, collected at the locations shown on the y-axis (see TABLE S1 for code explanation) and reared in the laboratory under 6-species gnotobiotic conditions.** CFU counts of AAB (red) and LAB (blue) recovered from homogenized pools of 2 flies each are shown. Data for 4-6 day old A,C,E,G,I,K) male and B,D,F,H,J,L) female flies moved from 25°C to A,B,G,H) 15°C, C,D,I,J) 25°C, or E,F,K,L) 32°C for 3 days immediately prior to homogenization are shown. Relative abundances are shown as the mean of AAB counts divided by the mean of LAB counts, with the fraction of LAB shown as a white point and the overlayed violin plot showing the distribution of fractional LAB abundance. Significant differences in relative abundances of LAB were determined by PERMANOVA (TABLE S4). Absolute CFU abundances are shown as the mean and standard error of the mean of all replicates. Significant differences in AAB and LAB abundance were determined by a Kruskal-Wallis test with a post-hoc Dunn test, and different letters over (AAB) or under (LAB) the bars report significant differences in their abundance

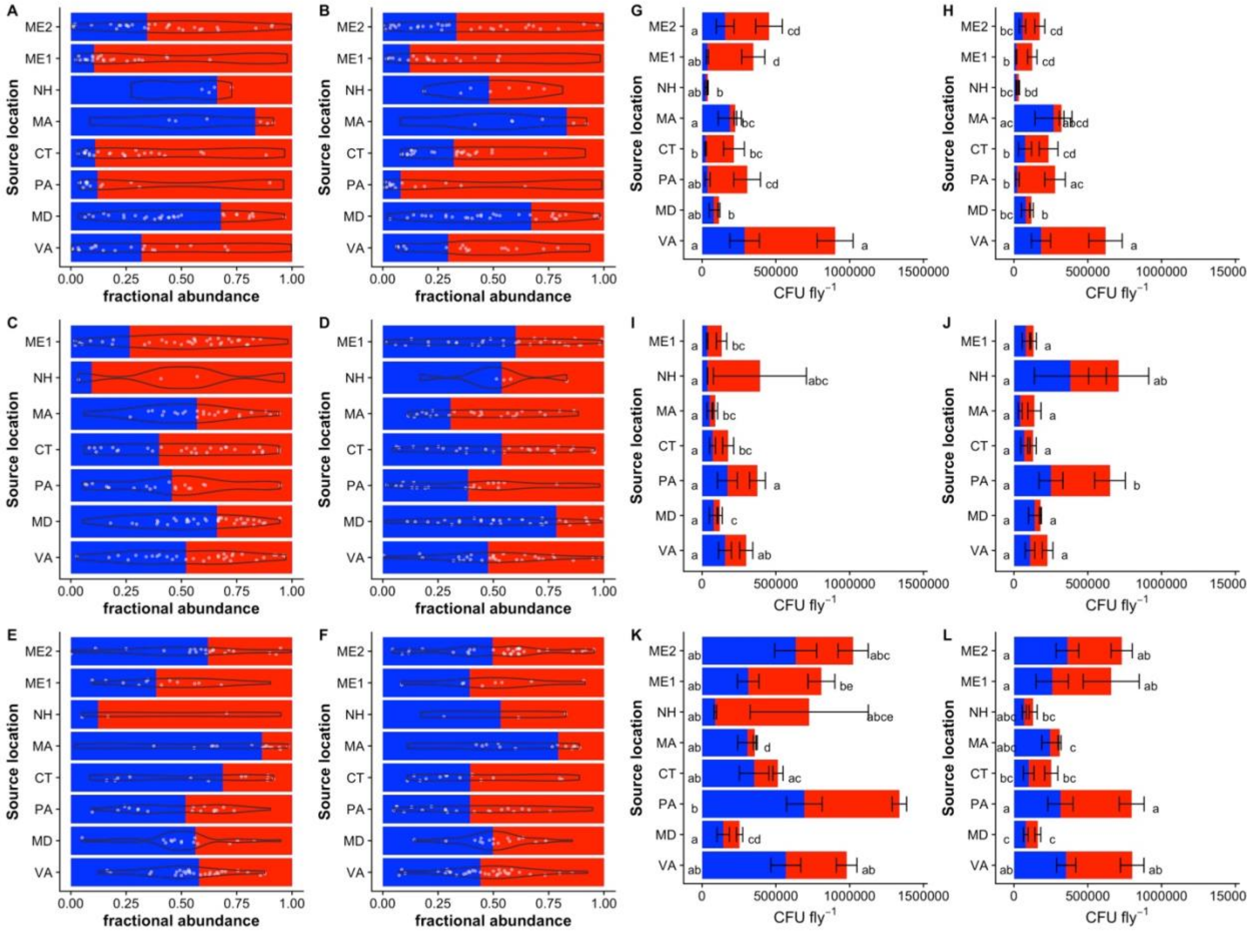

**Figure S4. Photoperiod-dependent variation in the microbiota composition of common garden fly populations, collected at the locations shown on the y-axis (see TABLE S1 for code explanation) and reared in the laboratory under 6-species gnotobiotic conditions.**

The relative A) and absolute B) abundances of AAB (red) and LAB (blue) colony forming units (CFUs) in flies reared at varying photoperiods are shown. Relative abundances are shown as the mean of AAB counts divided by the mean of LAB counts, with the fraction of LAB shown as a white point and the overlayed violin plot showing the distribution of fractional LAB abundance. Significant differences in relative abundances of LAB were determined by PERMANOVA (TABLE S4). Absolute CFU abundances are shown as the mean and standard error of the mean of all replicates. Significant differences in AAB and LAB abundance were determined by a Kruskal-Wallis test with a post-hoc Dunn test, and different letters over (AAB) or under (LAB) the bars report significant differences in their abundance.

The same data are shown, divided by sex and photoperiod condition, as CFU counts of AAB (red) and LAB (blue) recovered from homogenized pools of 2 flies each are shown. Data for 4-6 day old C,E,G,I,K,M) male and D,F,H,J,L,N) female flies moved from a 12h light:dark cycle to C,D,I,J) 1h light:23 h dark, E,F,K,L) 12h light:dark, or G,H,M,N) 23 h light: 1 hr dark cycle for 3 days immediately prior to homogenization are shown. Relative abundances are shown as the mean of AAB counts divided by the mean of LAB counts, with the fraction of LAB shown as a white point and the overlayed violin plot showing the distribution of fractional LAB abundance. Significant differences in relative abundances of LAB were determined by PERMANOVA (TABLE S4). Absolute CFU abundances are shown as the mean and standard error of the mean of all replicates. Significant differences in AAB and LAB abundance were determined by a Kruskal-Wallis test with a post-hoc Dunn test, and different letters over (AAB) or under (LAB) the bars report significant differences in their abundance

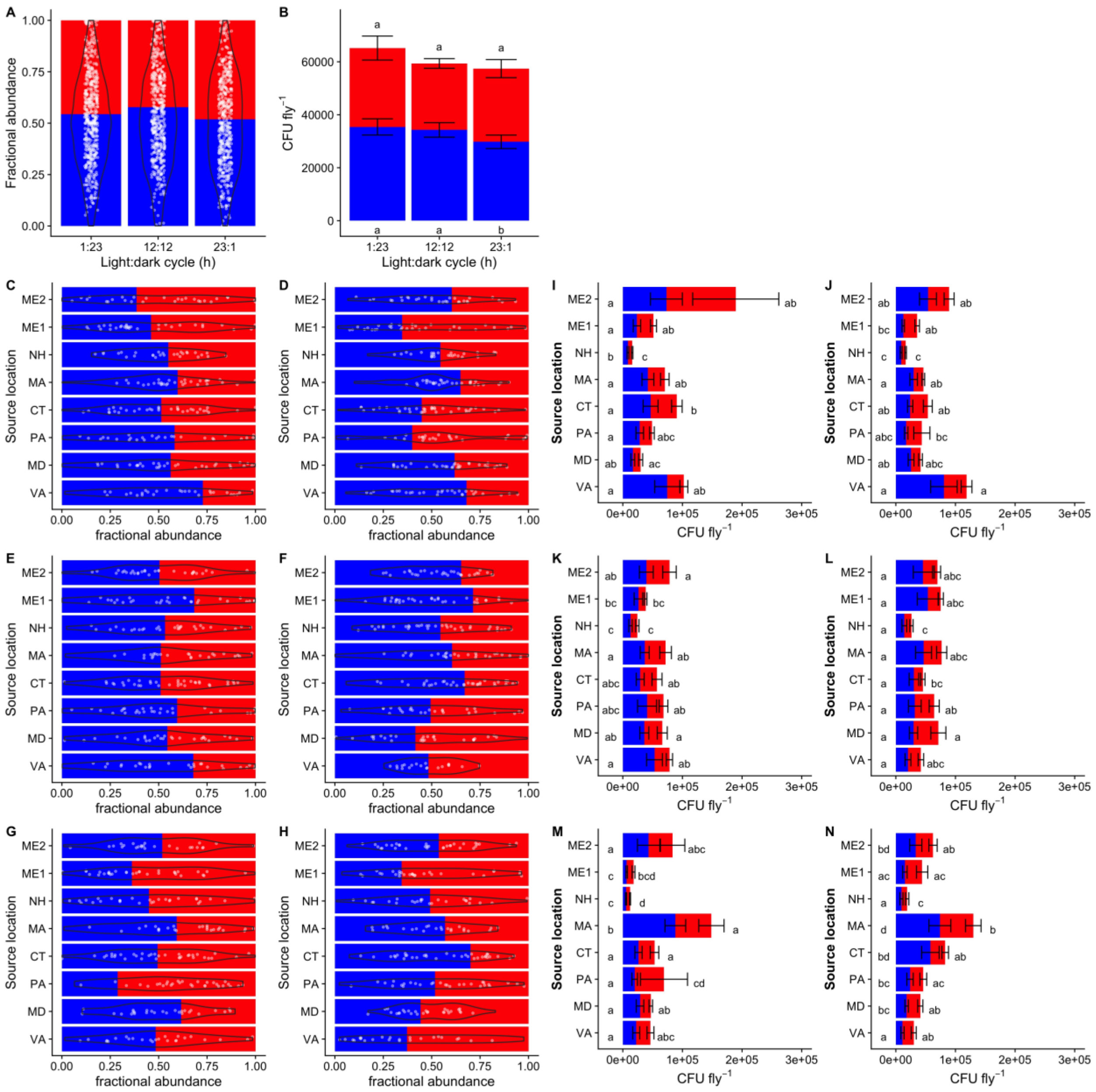

A
Fractional abundance
1.00
0.75
0.50
0.25
0.00
1:23
12:12
23:1
Light:dark cycle (h)
B
CFU fly$^{-1}$
60000
40000
20000
0
Light:dark cycle (h)
C
D
E
F
G
H
Source location
ME2
ME1
NH
MA
CT
PA
MD
VA
fractional abundance
I
J
K
L
M
N
0e+00
1e+05
2e+05
3e+05
CFU fly$^{-1}$

**Figure S5. Correlation between microbiota composition and time**. *Data correspond to Fig. 4B,I.* Correlation plots showing the data underlying Mantel tests that comparing the difference in time between samplings and the microbiota distance of the samplings from individual piles of peaches and apples established at different times over a fall season. Microbiota distance was defined using A-B) unweighted Unifrac and C-D) Bray Curtis distance metrics. The difference in time was calculated from A,C) calendar date (See Fig. 4B) or B,D) the time each pile was established. Black points show the distances between two samplings, and a trendline is shown in blue.

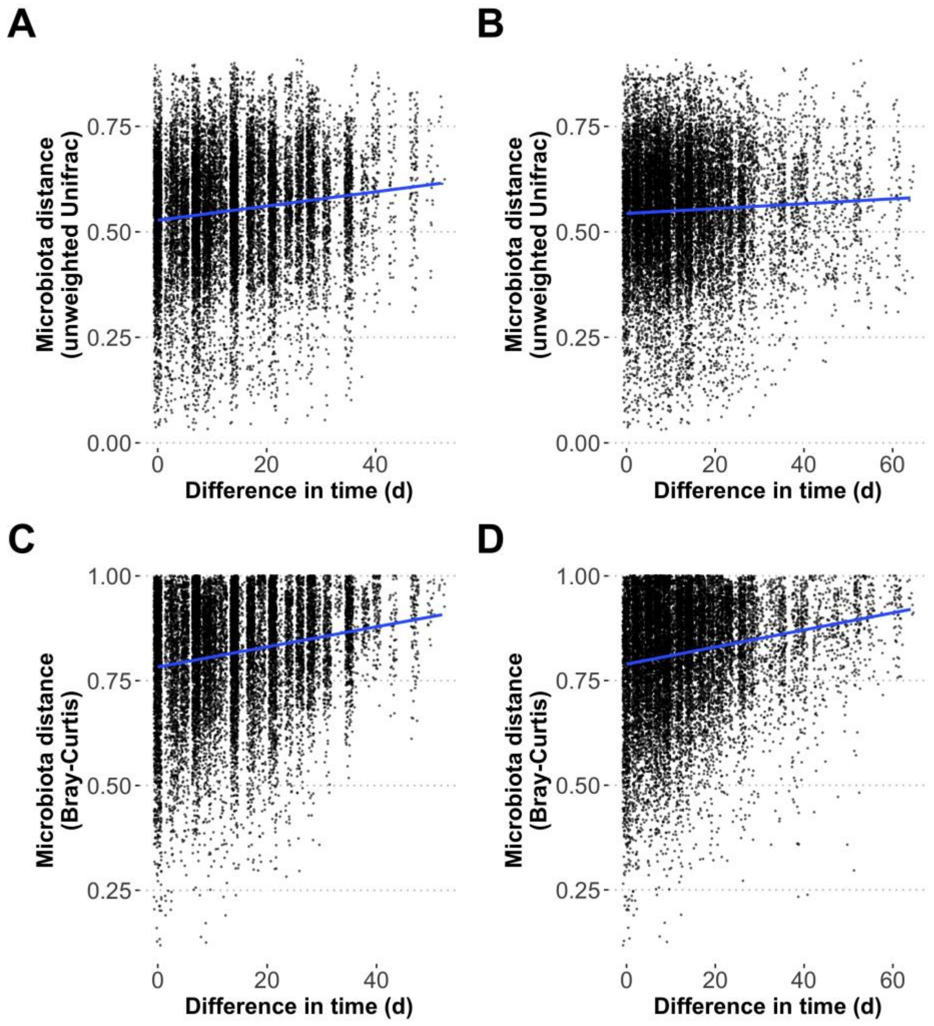

**Figure S6. Covariation between *Acetobacter* and *Commensalibacter* reads in different experiments.** *Data correspond to Fig. 1B-E and Fig. S1*. Samples were collected from the eastern USA in A) 2009 (from apples and peaches), B) 2018 (from grapes), C) 2018 (from apples), D) 2021 (from apples), or E) from Utah in 2020. Read counts were rarefied to 475 reads for A) and 500 reads for other studies. The Spearman's correlation coefficient and p-value for each comparison are shown, with a blue trendline through the data.

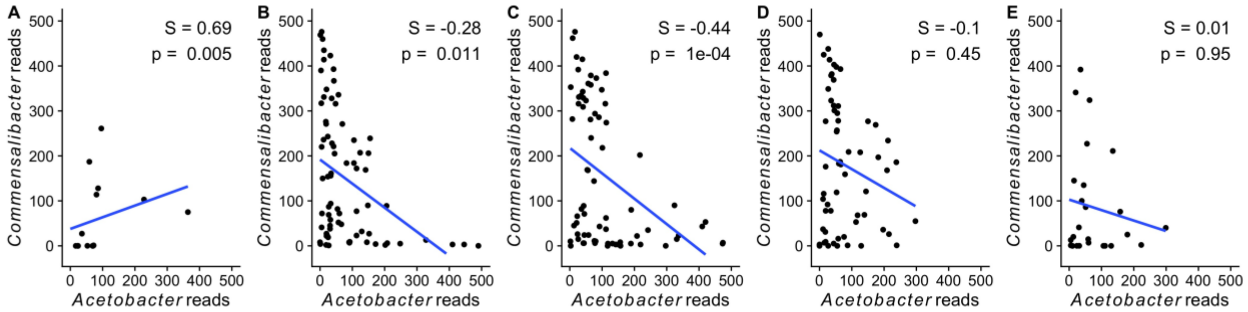

**Figure S7. Variation in microbiota composition with *Wolbachia* positivity in flies sampled from different fruits at an orchard in Middlefield, CT.** A) Taxon plot. ANCOM revealed significant differences in the abundance of B) *Acetobacteraceae* reads when clustered at the family level, or C) a specific *Commensalibacter* ASV when considered at the ASV level, between *Wolbachia*-negative and *Wolbachia*-positive flies

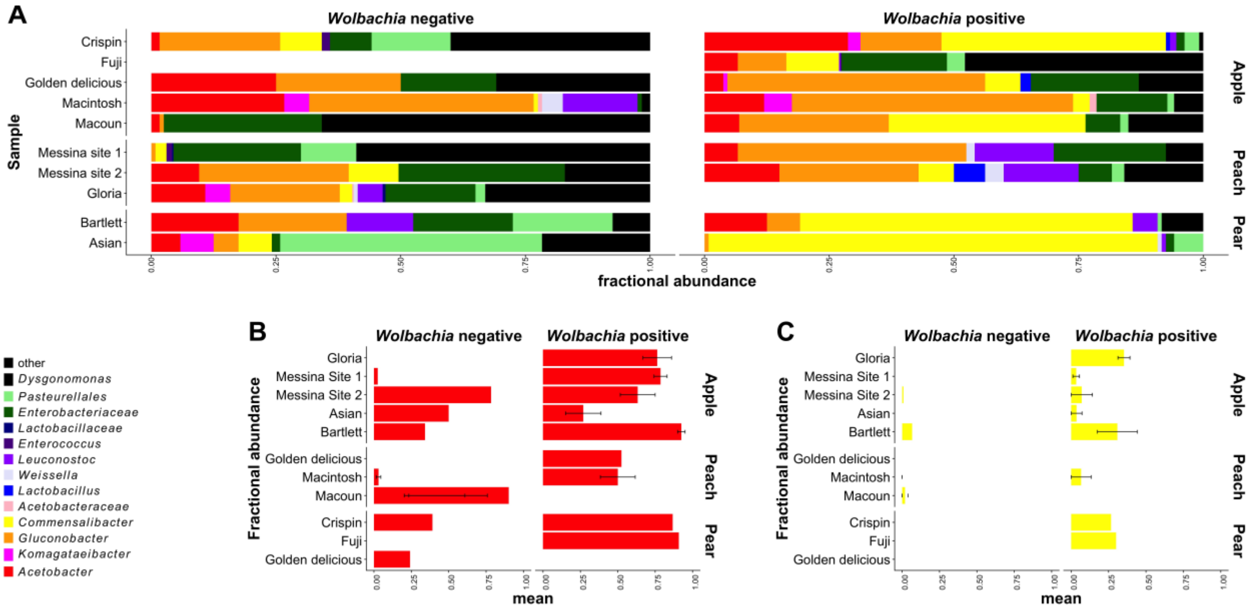

**Figure S8. Microbiota composition in gnotobiotic 6-species isofemale lines.** A-B) Male flies, C-D) female flies, A,C) absolute CFU counts, B,D) relative abundance. AAB CFUs (red) and LAB CFUs (blue) are shown. Different letters over or under the bars show significant differences by a Wilcoxon test (there are no differences). See Table S10 for PERMANOVA results that correspond to the data.

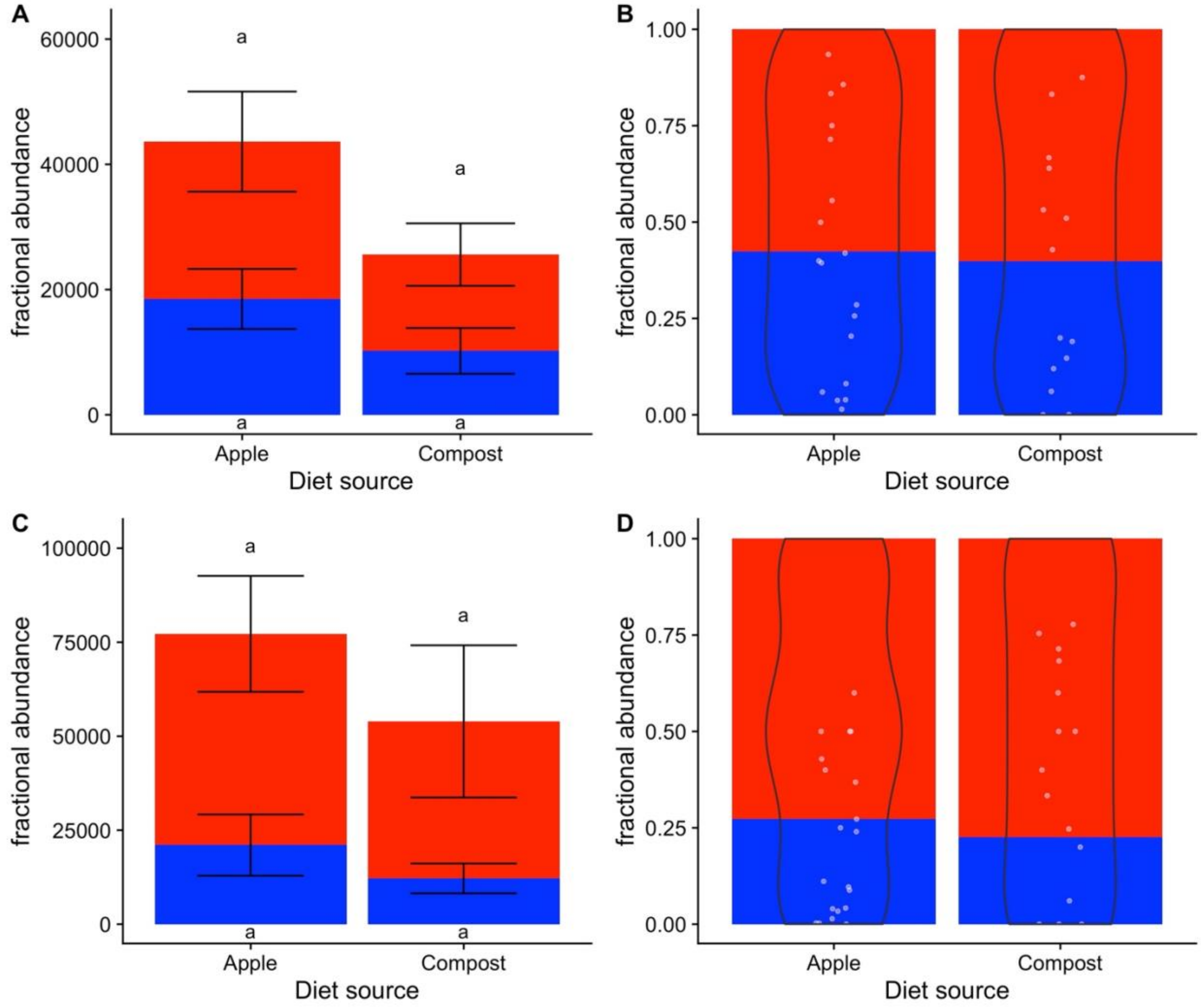
